# OXPHOS deficiencies affect peroxisome proliferation by downregulating genes controlled by the SNF1 signaling pathway

**DOI:** 10.1101/2021.08.23.457403

**Authors:** Jean-Claude Farré, Krypton Carolino, Lou Devanneaux, Suresh Subramani

## Abstract

How environmental cues influence peroxisome proliferation, particularly through other organelles, remains largely unknown. Yeast peroxisomes metabolize all fatty acids (FA), and methylotrophic yeasts also metabolize methanol. NADH and acetyl-CoA, the products of these pathways enter mitochondria for ATP production, and for anabolic reactions. During the metabolism of FA and/or methanol, the mitochondrial oxidative phosphorylation (OXPHOS) pathway accepts NADH for ATP production and maintains cellular redox balance. Remarkably, peroxisome proliferation in *Pichia pastoris* was abolished in NADH shuttling and OXPHOS mutants affecting complex I or III, or by the mitochondrial uncoupler, 2,4-dinitrophenol (DNP), indicating ATP depletion causes the phenotype. We show that mitochondrial OXPHOS deficiency inhibits the expression of several peroxisomal proteins implicated in FA and methanol metabolism, as well as in peroxisome division and proliferation. These genes are regulated by the Snf1 complex (SNF1), a pathway generally activated by high AMP and low ATP. Consistent with this mechanism, in OXPHOS mutants, Snf1 is activated by phosphorylation, but Gal83, its interacting subunit, fails to translocate to the nucleus. Phenotypic defects in peroxisome proliferation observed in the OXPHOS mutants, and phenocopied by the *Δgal83* mutant, were rescued by deletion of three transcriptional repressor genes (*MIG1*, *MIG2* and *NRG1*) controlled by SNF1 signaling. We uncovered here the mechanism by which peroxisomal and mitochondrial metabolites influence redox and energy metabolism, while also influencing peroxisome biogenesis and proliferation, thereby exemplifying interorganellar communication and interplay involving peroxisomes, mitochondria, cytosol and the nucleus. We discuss the physiological relevance of this work in view of human OXPHOS deficiencies.

## Introduction

Peroxisomes are ubiquitous organelles that are intimately involved in lipid metabolism [1, 2]. Like other subcellular organelles, they divide and segregate either to distribute themselves within cells or to endow daughter cells with peroxisomes. Cells stringently regulate organelle number, volume, size and content in response to environmental signals, which can emanate from other subcellular compartments. Regulation of these properties, together with organelle dynamics and homeostasis, allows cells to respond to metabolic or environmental stress, cope with the needs of cell division or differentiation, remove excess or damaged organelles by turnover, correct imbalances in organelle segregation during cell division or repopulate organelles with different enzymes upon switching to a new environment. The division of some organelles, such as the nucleus or the Golgi apparatus, is coupled to the cell cycle [3]. However, for others, such as mitochondria and chloroplasts, division is uncoupled from cell division [4]. Peroxisomes division can be coupled or uncoupled from cell division [5] and in yeast, peroxisome inheritance is cell-cycle regulated [6]. The de-coupling of peroxisome division during the cell-cycle and cell division is likely due to the ability of cells to also produce peroxisomes *de novo* [7].

In yeast and other eukaryotes, there are three major pathways that control peroxisome number. First, in constitutively-dividing cells, peroxisomes divide, as mitochondria and chloroplasts do, by fission of pre-existing peroxisomes, a process we refer to simply as peroxisome division [8, 9]. This process regulates peroxisome number in a geometric manner. A second general pathway is where peroxisomes are induced to create many new organelles within a short period of time, a process we call ‘peroxisome proliferation’. This increases organelle number much more rapidly than would be possible by sequential peroxisome division. Finally, peroxisome number is also controlled by pexophagy, the selective degradation of peroxisome by autophagic processes [10].

In yeasts, peroxisome division occurs both during constitutive growth and cell division. Peroxisome size and number are sensitive to peroxisomal metabolic pathways and the metabolites available [11–14]. Changes in peroxisome number and size can also be induced by peroxisome proliferation, which generally occurs when cells are shifted to nutrients whose metabolism requires peroxisomes and their constituent enzymes [1, 9]. Peroxisome volume changes in response to the import of matrix proteins [9]. Finally, peroxisomal contents can also vary, as shown for methylotrophic yeasts, in which peroxisomes are populated with enzymes involved in FA *β*-oxidation during growth in oleate, but new peroxisomes are endowed with methanol-assimilation enzymes upon growth in methanol [9].

The peroxisome division uses machinery also required for mitochondrial fission [15]. In yeast, this machinery is comprised of the proteins Fis1, Mdv and Caf4, as well as the dynamin-related GTPase, Dnm1 [16, 17]. The Pex11 family of proteins activates peroxisome division specifically [18–20], in a process that requires Pex11 phosphorylation to recruit Fis1 and Dnm1 to peroxisomes [21, 22]. A direct link to metabolite control of peroxisome division comes from *Yarrowia lipolytica*, where a signal from inside the peroxisomes (by the redistribution of acyl-CoA oxidase subunits from the peroxisome matrix to the membrane where they associate with the *Y. lipolytica* Pex16 protein) triggers this process [8]. Pex11 family proteins are conserved in evolution [15, 23] and are responsible for diseases in humans and plants [24–26]. In *Saccharomyces cerevisiae*, it has been proposed that while Pex11 promotes the division of peroxisomes already present in the cell, Pex25 initiates remodeling at the peroxisomal membrane to allow proliferation and Pex27 counters this activity [18].

Our current understanding of the mechanisms through which environmental cues are transduced into activation of peroxisome proliferation is limited [14, 23, 27, 28] and is largely unknown. In *S. cerevisiae*, peroxisome proliferation coincides with the transcriptional regulation by non-fermentable oleate of SNF1 complex-mediated signaling of several genes implicated in peroxisomal *β*-oxidation and peroxisome proliferation [29–32]. The Snf1 protein - the ortholog of the mammalian AMP-activated protein kinase (AMPK) - is a heterotrimer of the Snf1 catalytic subunit, the Snf4 activation subunit [33], and one of three *β* subunits (Sip1, Sip2, and Gal83) that localize the SNF1 complex to different cellular compartments [33, 34]. Gal83 directs the SNF1 complex to the nucleus in a glucose-regulated manner and facilitates the physical interaction between SNF1 and nuclear transcription factors [35]. In yeast cells grown in the absence of glucose, Snf1 is active via phosphorylation within its activation loop at Thr210 (T210) primarily by Sak1, but also by Tos3 and Elm1 [36–38]. The activated SNF1 complex inactivates the transcriptional repressors, Mig1 and Mig2 [39], by their phosphorylation [40, 41] and export from the nucleus, thereby enabling activation by the transcriptional activator, Adr1 [42]. There is no evidence of direct phosphorylation and activation of Adr1 by the SNF1 complex, but instead requires dephosphorylation of Adr1 at Ser230 by an unknown phosphatase [43]. Adr1 binds to upstream activating sequence 1 (UAS1) located adjacent to promoter sites during glucose-depression conditions.

Two additional transcriptional activators, Oaf1 and Pip2, regulate genes in response to oleate induction conditions [27, 31]. Oaf1 and Pip2 bind directly to oleate-responsive elements (OREs) in the promoter regions of coordinately-responsive peroxisomal matrix proteins and ORE binding sites are frequently found in proximity to UAS1 motifs [27]. However, these transcription activators have not been described in *P. pastoris*.

In contrast, the addition of glucose to cells (glucose repression) results in a reduction in ADP levels that causes the Glc7-Reg1 protein phosphatase to dephosphorylate T210 and thereby inactivate Snf1 [44–46].

Notably, this entire description of peroxisome induction and proliferation involves transactions of proteins and metabolites residing in three compartments – the cytosol, nucleus and peroxisomes, with no direct involvement of mitochondria. Yet, it is clear that peroxisomes have contacts with many subcellular compartments, including mitochondria [47–49]. Furthermore, mutations in genes affecting peroxisome biogenesis also impair mitochondrial function and morphology [50, 51], and the reverse is also true [52].

We decided to probe more deeply into the regulation of peroxisome content and proliferation in *P. pastoris*, which represents an excellent model as peroxisome content, number, size, and distribution change dramatically during growth in different carbon sources [14]. During growth in glucose, most *P. pastoris* cells possess a few, small peroxisomes, often next to the ER, with a limited lumenal content, but when cells are grown in oleate, numerous, small peroxisomes proliferate and are distributed throughout the cells, whereas in methanol medium the peroxisomes are large, less numerous, and clustered [53].

We reveal here the mechanism and players involved in the important interorganellar interplay involving the cytosol, nucleus, peroxisomes and mitochondria by which peroxisomal and mitochondrial metabolites influence redox and energy metabolism in the two compartments, while also influencing peroxisome biogenesis and proliferation.

## Materials and Methods

### Strains and plasmids are described in Supplemental Table S1

#### Media and reagents used to grow strains

All *P. pastoris* strains were prototrophic without requiring amino acid supplements or yeast extract and were cultivated in minimal media containing different carbon sources. The supplementation of media with amino acids did not change the results of this study.

YPD (2% glucose, 2% bacto-peptone, 1% yeast extract), glucose medium (2x YNB [YNB: 0.17% yeast nitrogen base without amino acids and ammonium sulfate, 0.5% ammonium sulfate], 0.04 mg/L biotin, 2% dextrose), oleate medium (2x YNB, 0.04 mg/L biotin, 0.02% Tween-40, 0.2% oleate), and methanol medium (2x YNB, 0.04 mg/L biotin, 1% methanol). Amino acids were supplemented when indicated in the results section using 0.79 g/L complete synthetic medium of amino acids and supplements (CSM; #1001, Sunrise Science Products, USA). 2,4-Dinitrophenol (DNP; #D198501, Sigma-Aldrich, USA) was dissolved in methanol, used at a final concentration of 0.25 mM and added after switching from glucose to the methanol media. The total concentration of methanol in the culture containing DNP was 1%.

#### Plasmid constructions

Plasmids were constructed by Gibson Assembly for which primers were designed using NEBuilder (https://nebuilder.neb.com/#!/). DNA was amplified from WT genomic DNA by PCR using Advantage 2 Polymerase (#639202, Takara Bio, USA). Plasmid backbones were double-digested with the necessary restriction enzymes and purified using the Qiagen DNA purification kit (#28704X4, QIAGEN GmbH, Germany). DNA inserts were cloned into the digested vectors using the NEBuilder Hifi DNA Assembly Mastermix (#E2621L, New England Biolabs) and incubated for 1h at 50° C.

#### Yeast strain constructions

Plasmids were linearized with the appropriate restriction enzyme before transforming by electroporation into competent yeast cells, as previously described [54]. The transformed cells were plated on YPD plates with the appropriate yeast selection markers and incubated at 30° C for a few days. Colonies were screened by Western Blot or fluorescence microscopy.

#### Fluorescence microscopy

Cells were grown in YPD at 30° C until exponential phase (1-2 OD_600_/ml), washed twice with sterile water, and then transferred to glucose media or peroxisome proliferation (methanol or oleate) media. With the goal of keeping cells at exponential phase, different starting OD_600_ were used for different strains, depending on the strain’s cell doubling time in the respective media. Cells were grown in indicated media in 250 mL flask at 30° C and shaken at 250 rpm. Cells were pelleted, washed twice with sterile water and 1.5μl of cells were mixed with melt 1% low melting point agarose and placed on a glass slide with a cover slip and imaged using 63x or 100× magnification on a Carl Zeiss Axioskop fluorescence microscope. Images were taken on an AxioCam HR digital camera, no digital gain was used, exposition for peroxisome markers were kept constant during microscopy of different strains. Images were processed on AxioVision software. The images presented are representative results from experiments conducted at least in triplicate.

#### Biochemical studies

Same condition described for fluorescence microscopy assays were used to grow cells. Five OD_600_ of cells was collected at different times as described in the figures, trichloroacetic acid precipitated and analyzed by Western blot.

#### Mitoplate assay

Mitochondrial metabolic activity was measured in digitonin-permeabilized cells using the PM1 MicroPlate (Biolog #12111, Hayward, CA, USA) following the manufacturer’s instructions (protocol document dated February 6, 2019 using cell preparation protocol Option 1).

Briefly, cells were grown in lactate/glycerol media, washed and resuspended in high lactose osmotic stabilizing solution (YMAS), containing digitonin (#D-180-250, Gold Biotechnology, USA). After 1 h, the permeabilized cells were seeded into 96-well PM1 MicroPlates containing different potential energy substrates. The colorimetric assay was initiated by adding Redox Dye Mix MC (Biolog #74353). The PM1 MicroPlate was then loaded into the OmniLog PM-M system (Biolog, Hayward, CA, USA) for kinetic reading. Assays were performed in triplicate.

*qRT-PCR* - RNA extraction was performed using Trizol™ Reagent (#15596026, Invitrogen, USA) following the manufacturer’s instructions with some modifications. Twenty OD_600_ of frozen cells, grown in the conditions described in the results section, were resuspended with 1 mL of Trizol and 250 μl of acid-washed glass beads (425-600 microns). The suspension was vortexed for 30 sec, followed by chilling on ice for 30 sec. This step was repeated three times. The extraction was completed according to the protocol described by the kit manufacturer and was followed by DNAse I treatment (#18068015, Life technologies, Carlsbad, CA, USA). The RNA quantity and quality were determined using a DU spectrophotometer (Beckman Coulter Inc., Fullerton, USA) to determine the 260/280 nm ratio. The RNA extractions showed a 260/280 ratio of approximately 1.8–2.0, confirming good RNA quality. The RNA samples were stored at −80° C. cDNA was synthesized from 1 μg of total RNA using an Invitrogen two-step kit with SuperScript™ III (#18080051, Invitrogen, Carlsbad, CA, USA) as the reverse transcriptase (RT) enzyme with random hexamer. This was followed by RNAse treatment, according to the manufacturer’s instructions. The cDNAs were stored at −80° C. One μl of cDNA (equivalent to 80 ng of total RNA), 300 nM of primers listed in Supplementary Table S2 and PowerUp™ SYBR™ Green master mix (#A25741, Applied Biosystems, USA) were used following the manufacturer’s instructions. Values for each target gene were normalized using 18S rRNA and WT in glucose condition as reference. Expression values were calculated using the 2^−△△CT^ method [55].

## Results

### Peroxisome metabolites influence peroxisome size

We reinvestigated whether peroxisomal metabolites could influence peroxisome size and number in *P. pastoris*. Peroxisomes were labelled with the functional, fluorescently-tagged, peroxisomal membrane proteins (PMPs), Pex3-GFP or GFP-Pex36, in several mutant strains grown in different carbon sources (Fig. 1A and Fig. S1A). We then deleted key genes encoding enzymes of the FA *β*-oxidation and methanol metabolism pathways, as well as a peroxin required for the import of peroxisome matrix proteins. Supporting previous findings, wherein intermediates of peroxisome metabolism, or the absence of certain peroxisomal enzymes, regulate the maturation and fission of the organelle [56, 57], the lack of 3-ketoacylCoA thiolase (Pot1), an enzyme responsible for the last step of FA *β*-oxidation, also affected peroxisome proliferation exclusively in oleate (Fig. 1A, S1A). Similarly, the lack of alcohol oxidase (Aox) 1 and 2, the first enzymes in the methanol utilization pathway (MUT), affected peroxisome proliferation only in cells grown in methanol medium (Fig. S1A). In the same way, the lack of the peroxin, Pex5, responsible for the import of PTS1-containing enzymes (including some involved in the *β*-oxidation and MUT pathways), impaired peroxisome proliferation in both media (Fig. S1B). These results show that one or more peroxisomal intermediate metabolite/s induces peroxisome proliferation, in addition to previous reports regarding their effects on division.

**Figure 1:**
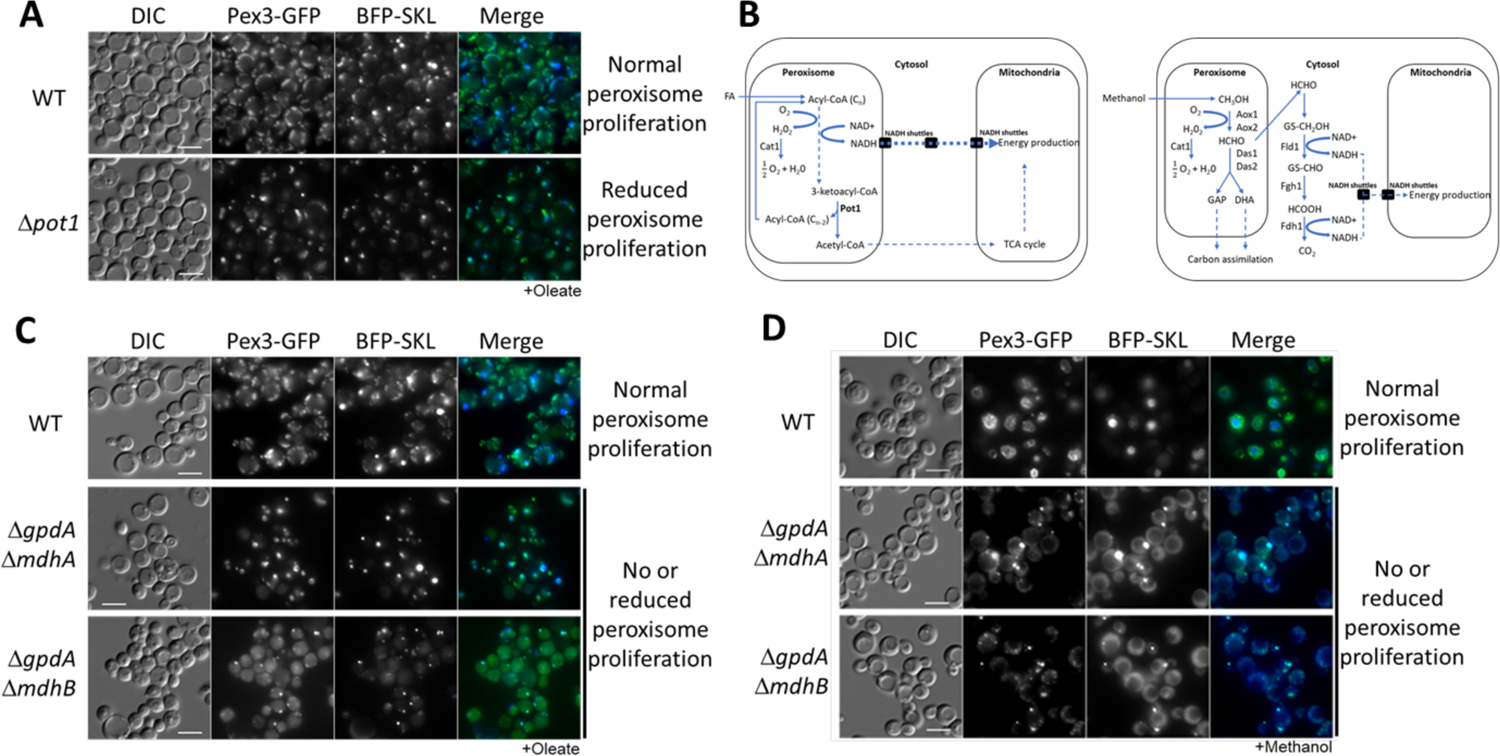
Peroxisome metabolites influence peroxisome size. (A) Fluorescence microscopy of WT and *Δpot1* mutant cells expressing Pex3-GFP driven by *PEX3* promoter and BFP-SKL driven by the *GAPDH* promoter, grown in oleate for 8 h. (B) Brief description of FA *β*-oxidation, methanol metabolism and NADH shuttling between peroxisomes and mitochondria. Cat1, catalase; Pot1, 3-ketoacylCoA thiolase; Aox, alcohol oxidase; Das, dihydroxyacetone synthase; Fld1, formaldehyde dehydrogenase; Fgh1, S-formylglutathione hydrolase; Fdh1, formate dehydrogenase. (C) and (D) Fluorescence microscopy of WT and NADH shuttling mutant cells expressing Pex3-GFP driven by the *PEX3* promoter and BFP-SKL driven by the *GAPDH* promoter, grown in oleate and methanol for 8 h, respectively. Bars: 5 μm.

Although some of the transcriptional networks in the induction of peroxisomes are known, the molecular target of the signaling pathway that triggers peroxisome proliferation has not yet been elucidated, so we reevaluated the concept. We hypothesized that if the molecule(s) inducing proliferation localize(s) in the peroxisome lumen, it would need to communicate with the cytosol and/or other organelles to induce the lipid and membrane transfer needed for peroxisome growth.

Two mechanisms have been considered for such lipid and/or membrane transfer - through ER-derived vesicles that fuse with immature peroxisomes [58] and/or by non-vesicular lipid transfer originating at interorganelle membrane contact sites [59]. Additionally, there must be activation of the fission machinery needed to increase peroxisome numbers (including peroxisomal division factors like Pex11, which recruits to the peroxisomes shared components of the mitochondrial fission machinery [18]).

In *S. cerevisiae*, the final intraperoxisomal products of *β*-oxidation are NADH and acetyl-CoA, for cells grown in FA (Fig. 1B, [2]). The transport of reducing equivalents out of peroxisomes and the maintenance of the intraperoxisomal redox balance in *S. cerevisiae* is mediated by the malate/oxaloacetate shuttle during growth in oleate, and by both the malate/oxaloacetate and the glycerol-3-phosphate/dihydroxyacetone phosphate (G3P/DHAP) shuttles, during growth in glucose [60]. The malate/oxaloacetate shuttle is coordinated by three NAD^+^-dependent malate dehydrogenases: the mitochondrial Mdh1 that is part of the TCA cycle, the peroxisomal Mdh3 which regenerates NAD^+^ for the FA β-oxidation process, and the cytosolic Mdh2 probably involved in the glyoxylate cycle [61]. The G3P/DHAP shuttle is coordinated by two G3P dehydrogenases: the mitochondrial Gpd2 and peroxisomal/cytosolic Gpd1 [62, 63]. *P. pastoris* possesses only two malate dehydrogenases (designated MdhA or B) and one G3P dehydrogenase (GpdA). We recently characterized these enzymes and reported that MdhA and GpdA are predominantly mitochondrial enzymes in all media analyzed, whereas MdhB is cytosolic during growth in glucose or methanol, but is cytosolic/peroxisomal during growth in oleate [64].

With the intent of blocking the shuttling of NADH to the mitochondria, we made single and double deletions of both NADH shuttles in *P. pastoris* and we analyzed peroxisome status after 24 h of induction in oleate medium, by fluorescence microscopy, using Pex3-GFP and BFP appended with a PTS1 (BFP-SKL) (Fig. 1C, 1D and Fig. S2). We did not observe any peroxisome defect in the *gpdA* mutant (Fig. S2), consistent with its NADH-shuttling role in glucose, but not in oleate medium, in *S. cerevisiae* [60]. However, both the *mdhA* and *mdhB* mutants had partial defects in peroxisomal division and/or maturation (Fig. S2). In the *mdhA* mutant, peroxisome numbers were strongly reduced, and we observed peroxisome tubulation characteristic of *Δfis1* cells. In contrast, the *mdhB* deletion exhibited a mixed phenotype, wherein some cells seemed to contain only a single, tiny, import-competent peroxisome, but some other cells showed several import-competent peroxisomes, although they remained more clustered than in wild-type (WT) cells (Fig. S2). Interestingly, both double deletion strains, *ΔgpdA ΔmdhA* and *ΔgpdA ΔmdhB* cells, showed a more consistent phenotype, with most of the cells containing only a single, import-competent peroxisome, when cells were grown in oleate (Fig. 1C). These results are consistent with the role of NADH-shuttling proteins in maintaining the intraperoxisomal redox balance and peroxisome proliferation during growth in oleate, but might also indicate that the G3P/DHAP shuttle contributes to the redox balance during *β*-oxidation of FA in *P. pastoris*.

Peroxisomes metabolize methanol to formaldehyde, which diffuses to the cytosol and is oxidized by a dehydrogenase (Fld1) to formate, which is further oxidized by a second dehydrogenase (Fdh1) to carbon dioxide, yielding NADH (Fig. 1B). This NADH then shuttles to the mitochondria to maintain the cytosolic redox balance and to feed mitochondrial OXPHOS for energy production. The phenotypes of the double deletion strains were relevant because during growth in methanol as the only carbon source, the redox reactions needed from methanol metabolism occur in the cytosol, and intraperoxisomal redox should not be altered. Consequently, we expected the signal driving peroxisome to be present to allow peroxisome proliferation.

However, surprisingly, the same double deletions strains showed a similar phenotype of reduced peroxisome proliferation, when cells were grown in methanol medium for 24 h (Fig. 1D), suggesting an extra-peroxisomal proliferation signal in this case.

### Mitochondrial OXPHOS mutants also affect multiple peroxisome phenotypes

We hypothesized that NADH, which shuttles from the peroxisomes to the mitochondria during growth in oleate medium, and from the cytosol to the mitochondria during growth in methanol medium, maintains the redox balance that sustains peroxisome metabolism but also contributes to energy production and could be the molecule triggering peroxisome maturation and proliferation. In *P. pastoris*, during growth in methanol or oleate media, NADH shuttles to the mitochondria where it is oxidized by the respiratory chain. The NADH dehydrogenase complex I (CI) is the main, energy-conserving, entry point for electrons into the respiratory chain. This enzyme directly oxidizes NADH to NAD^+^ and concomitantly reduces coenzyme Q (CoQ, ubiquinone), which causes H^+^ to be pumped out across the inner mitochondrial membrane, to contribute to the proton gradient that drives ATP synthesis [65].

To study the involvement of mitochondrial NADH oxidation in peroxisome proliferation and fission, we deleted two, nuclear-encoded genes encoding the mitochondrial complex I, the accessory subunit, Ndufa9, and the core subunit, NugM [65]. Because none of these mutants have been previously characterized in *P. pastoris*, we assessed their necessity for the electron transport chain in response to metabolic substrates using Biolog’s Yeast Mitochondrial Energy Substrate Assays. The permeabilized WT or mutant cells, pre-grown in glycerol/lactate media, were seeded into 96-well plates containing diverse cytoplasmic and mitochondrial metabolic substrates, and incubated at 30° C for 16 h. The metabolism of substrates was assessed by monitoring the colorimetric change of a terminal electron acceptor, a tetrazolium redox dye. As expected, mitochondrial substrates, producing NADH for Complex I such as L-lactic acid, citric acid, *α*-keto-glutaric acid, pyruvic acid, and others were properly metabolized in WT, but not in the mutant cells (Fig. 2A). In sharp contrast, Complex II substrates, *α*-glycerol-phosphate, succinic acid and methyl succinate, exhibited unchanged rates of metabolism in the mutants. This result also suggests that Complex III activity was not impaired in these Complex I mutants.

**Figure 2:**
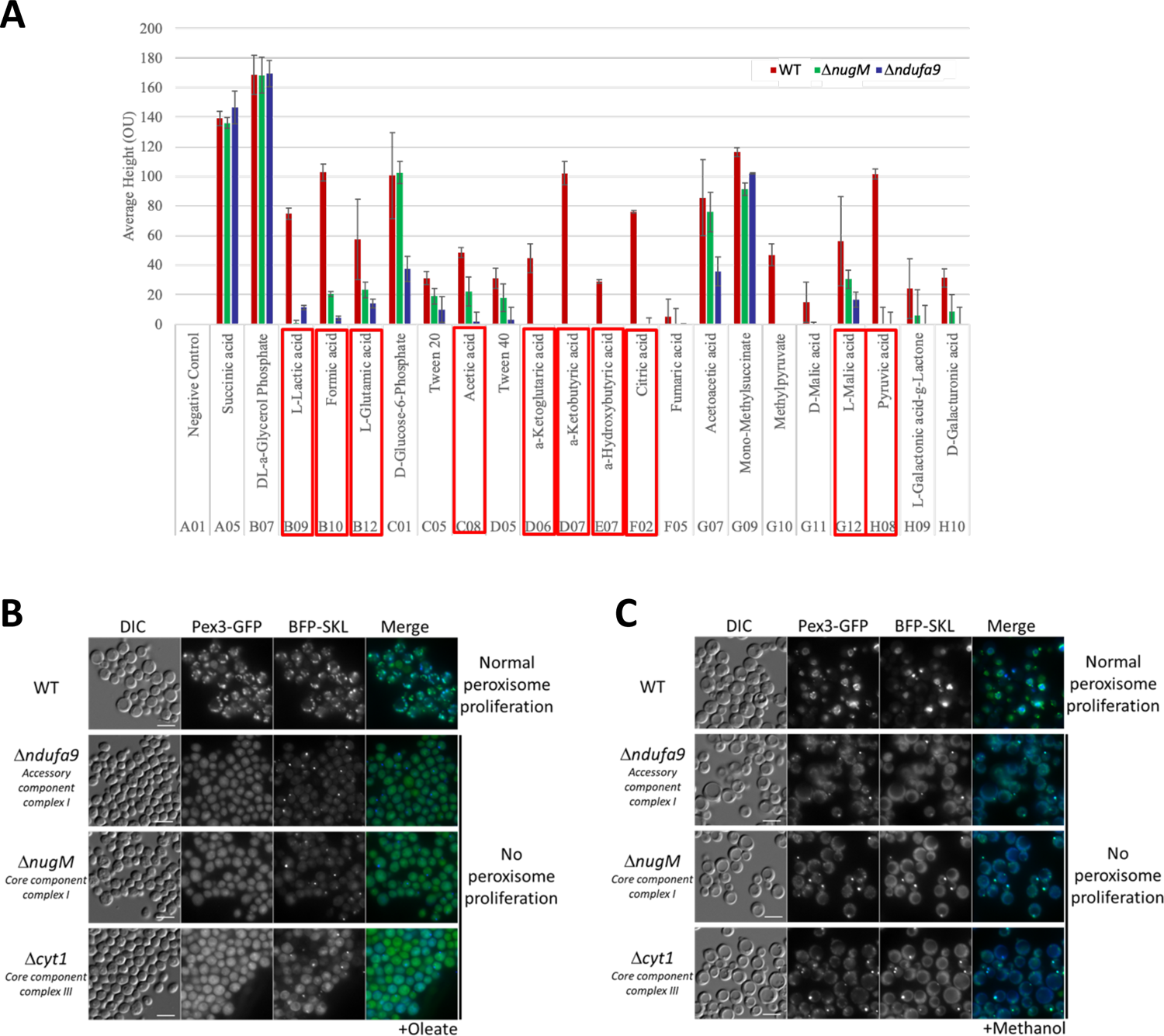
Altered mitochondrial respiration in cells with CI mutants and dysfunctional mitochondria affect peroxisome proliferation. Equal amounts of WT, *Δndufa9* and *ΔnugM* cells were grown as described in Materials and Methods, and mitochondrial metabolic activity was measured in digitonin-permeabilized cells using the PM1 MicroPlates from Biolog Inc. Results are presented as the average rate/min/μg of protein (OU). Respiratory substrates are highlighted with boxes. (B, C) Fluorescence microscopy of WT, *Δndufa9*, *ΔnugM* and *Δcyt1* mutant cells expressing Pex3-GFP driven by the *PEX3* promoter and BFP-SKL driven by the *GAPDH* promoter, grown for 8 h in (B) oleate and (C) methanol medium. Bars: 5 μm.

The peroxisome status in the CI mutants was monitored by fluorescence microscopy after overnight growth in oleate and methanol media (Fig. 2B, C). Remarkably, unlike the situation in WT cells, we did not observe an increase in peroxisome number or size over time in both CI mutants in both media, nor was there any peroxisome proliferation. While both CI mutants were competent for the peroxisomal import of BFP-SKL, most cells had only a single peroxisome that failed to proliferate in oleate or methanol, or divide.

Due to the direct role of CI in NADH oxidation, we hypothesized that it might participate directly in peroxisome biogenesis and somehow transmit a signal to activate the proliferation. To differentiate a direct role in peroxisome proliferation from the general role of electron transfer and proton translocation by CI, we deleted a gene encoding a downstream component of the OXPHOS complex, but not directly implicated in NADH oxidation, the *CYT1* gene, encoding a core subunit of complex III. As seen earlier for the CI mutants, the *Δcyt1* mutant also blocked peroxisome proliferation (Fig. 2B, C). This result indicates that peroxisome proliferation depends on a fully functional mitochondrial respiratory chain.

The similarity in peroxisome proliferation defects between mutants of CI and CIII suggested that the lack of ATP synthesis, and not the redox balance, is the cause of these phenotypes. To verify this hypothesis, we used the mitochondrial uncoupler, DNP, which separates the flow of electrons from the pumping of H^+^ ions for ATP synthesis [66]. DNP transports protons across the mitochondrial inner membrane, altering the proton gradient and inhibiting ATP production via OXPHOS. We followed peroxisome proliferation in methanol after 3 and 24 h, using WT cells expressing Pex3-GFP driven from its own promoter (Fig. 3). DNP fully abolished peroxisome proliferation and only a single, small peroxisome was observed in each cell, confirming our hypothesis that ATP synthesis is necessary in signaling peroxisome proliferation.

**Figure 3:**
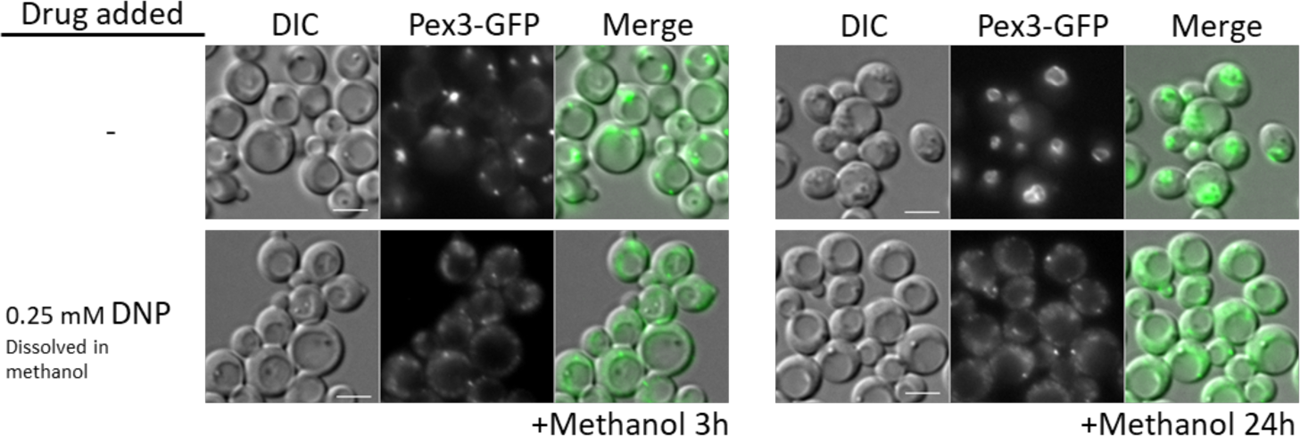
WT cells treated with the OXPHOS uncoupler, DNP, share peroxisome proliferation defects with mitochondrial CI and CIII mutant cells. Fluorescence microscopy of WT cells expressing Pex3-GFP driven by the *PEX3* promoter, grown in methanol medium, with or without 0.25 mM DNP. Bars: 5 μm.

### The OXPHOS effect on peroxisome proliferation acts via the Snf1 kinase pathway in *P. pastoris*

To understand the consequences of OXPHOS deficiency on peroxisome proliferation, we checked the relative protein abundance of the key peroxisome division factor, Pex11 (Pex11-2HA expressed from its own promoter, P*_PEX11_*) after 4 h of induction in oleate medium (Fig. 4A). Remarkably, we did not detect Pex11-2HA in *ΔnugM* cells, suggesting that an inactive SNF1 pathway could account for the OXPHOS deficiency. In *S. cerevisiae*, where the regulation of peroxisome proliferation by the Snf1 pathways is the best understood, the proliferative capacity of peroxisomes coincides with the FA-responsive transcriptional regulation of many genes encoding peroxisomal proteins [67]. Many such genes are repressed in glucose [68, 69] and derepressed in oleic acid [70]. The derepression of these genes upon removal of glucose depends on the SNF1 signaling pathway. As stated in the Introduction, the SNF1 complex achieves this regulation via the interplay of several transcriptional regulators, such as the transcriptional activator, Adr1[42], and the transcriptional repressors, Mig1 and Mig2 [39–41].

**Figure 4:**
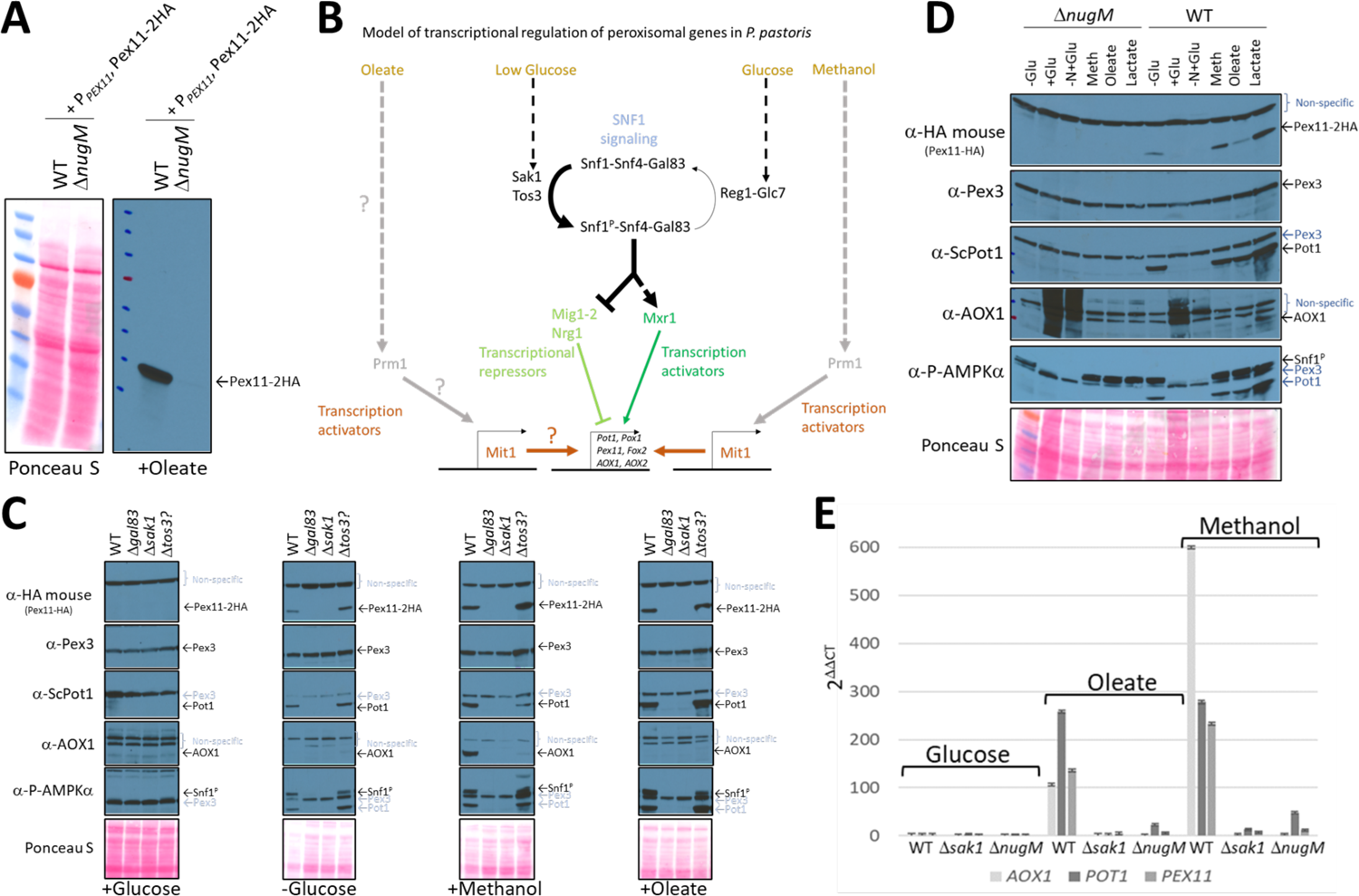
OXPHOS mutants, like SNF1 mutants impair Pot1, Aox1 and Pex11 expression. (A) Western blot of Pex11-2HA visualized with anti-HA antibodies in WT and *ΔnugM* mutant cells. (B) Model of transcriptional regulation of peroxisome genes regulated by SNF1 signaling.denotes unknown pathway for induction of Mit1 in oleate. (C) and (D) western blots of several peroxisomal proteins and the phosphorylated form of Snf1 in WT, SNF1 complex mutants and/or *ΔnugM* mutant cells. Non-specific bands and signal arising from previous western blots are indicated in blue font. Ponceau S staining was used as a loading control. (E) Relative expression of *PEX11*, *POT1* and *AOX1* was obtained after 4 h incubation in the indicated carbon source using 18S ribosomal RNA (18S-rRNA) for qPCR normalization, and WT in glucose medium as reference and the 2^−ΔΔCT^ method for the analysis [112].

In *P. pastoris,* the regulation of peroxisomal genes involved in *β*-oxidation has not been studied much. However, due to the strong interest in *P. pastoris* as a versatile cell factory for the production of recombinant proteins both on laboratory and industrial scales, many studies have focused on the regulation of the *AOX1* promoter (P*AOX1*) during growth in glucose, glycerol and methanol media [71–74]. Similar to *β*-oxidation genes, *AOX1* is repressed during growth in glucose and strongly induced by methanol. The transcriptional activator, Mxr1 (which shares sequence and functional homology with *S. cerevisiae* Adr1) [74, 75] and the transcriptional repressors, Mig1, Mig2 and Nrg1 [71, 76], (involved in glucose repression) regulate the derepression from glucose in *P. pastoris* (Fig. 4B). As in *S. cerevisiae*, these transcription factors are most probably regulated by the SNF1 pathway, as a recent high-throughput screen implicated Sak1, the primary Snf1-activating kinase, and Gal83, the *β*-subunit of the SNF1 complex, in the expression of AOX [77]. In addition, Mxr1 is inactivated by a phosphorylation at Ser 215 (S215) [75].

In *P. pastoris,* the *S. cerevisiae* transcription activators, Pip2 and Oaf1, have not been found by *in silico* studies and it has been suggested they are absent in methylotrophic yeast. But methanol and glycerol activate at least two transcription factors, Mit1 and Prm1, which induce enzymes required for methanol utilization (MUT), but not for peroxisome proliferation, by a mechanism resembling Pip2-Oaf1 activation [73, 78]. During methanol induction, Prm1 transmits the methanol signal to Mit1 by binding to the *MIT1* promoter, thus increasingly expressing Mit1 and subsequently activating the promoter of methanol metabolism genes, such as *AOX1* [73].

We analyzed Pex11-2HA, Aox1, Pot1 and Pex3 expression levels under different carbon induction conditions using available kinase deletions strains affecting the SNF1 signaling pathway (Fig. 4C) [77]. As expected, in WT cells, Pex11-2HA, Pot1 and Aox1 were not detected in glucose medium, but glucose derepression (-Glucose) was enough to induce Pex11-2HA and Pot1, but not Aox1, which has no role during the growth of WT *P. pastoris* in oleate. Glucose derepression, when combined with the addition of methanol, was needed to observe Aox1 expression in WT cells in our cultivation conditions, indicating a more complex regulation, such as the activation by methanol of the transcription activators, Mit1 and Prm1, in agreement with previous studies on regulation of the *AOX1* promoter [73].

We examined the activation of Snf1 by phosphorylation next because glucose derepression acts via the SNF1 pathway. The amino acid sequence near the activation loop of *P. pastoris* Snf1 (including Thr 171) is identical with that in human AMPKα [79]. Thus, a phospho-AMPKα (Thr 172) antibody designed to correspond to the residues surrounding Thr 172 of human AMPKα should detect the active form of *P. pastoris* Snf1. As seen in other organisms*, P. pastoris* Snf1 was activated in WT cells in response to glucose limitation by phosphorylation of Thr 171 in the activation loop of its catalytic subunit (Fig. 4C). A mutant of a putative Tos3 kinase (Tos3?; UniProt gene name: PAS_chr1-3_0213), with weak homology to its *S. cerevisiae* counterpart, from the *P. pastoris* kinase deletion collection was not required for Snf1 activation, nor for the downstream SNF1 regulation, indicating that PAS_chr1-3_0213 is not required for SNF1 signaling or for peroxisomal protein expression. In contrast, Gal83 and Sak1 were essential for Snf1 phosphorylation and the expression of Pex11-2HA, Pot1 and Aox1 in every condition tested, confirming their major role in the SNF1 signaling pathway for the expression of some peroxisomal proteins. We did not observe regulation of the peroxisome biogenesis factors, Pex3 or Pex17, by the SNF1 pathway or by the carbon sources in our incubation conditions.

We also analyzed Snf1 activation and expression of the same peroxisomal proteins in the *ΔnugM* strain using the same experimental conditions, and remarkably we observed similar expression defects as those observed for *Δsak1* and *Δgal83* strains, despite phosphorylation of Snf1 (Fig. 4D). We previously earlier that the mitochondrial uncoupler, DNP, causes a similar peroxisome proliferation defect as the OXPHOS mutants, and like the *ΔnugM* mutant also affects the expression of Pex11, Pot1 and Aox1 during methanol cultivation, despite phosphorylation of Snf1 (Fig. S3A). Thus, DNP phenocopies the OXPHOS mutants in this respect.

If dysfunctional mitochondria impair SNF1 signaling directly, peroxisomal mRNA levels of *PEX11*, *POT1* and *AOX1* should be downregulated, like what is observed for an inactive SNF1 complex in *S. cerevisiae* [32, 70]. We performed quantitative real-time RT-PCR (qRT-PCR) analysis and showed that all these mRNAs were upregulated in WT during glucose derepression and/or peroxisome proliferation conditions, as expected. In WT cells, the mRNA levels of *PEX11* and *POT1* were upregulated after glucose removal and during cultivation in oleate or methanol medium, while *AOX1* mRNA upregulation was mostly observed in methanol medium (Fig. 4E). However, confirming a potential role of OXPHOS in SNF1 signaling, *ΔnugM* cells, like the *Δsak1* cells, were unresponsive to glucose derepression and/or oleate and methanol cultivation, and mRNA levels were not significant upregulated, relative to levels seen in WT cells, in any of the conditions tested.

### OXPHOS defect in peroxisome proliferation is not rescued by activation of known transcriptional activators of genes encoding peroxisomal proteins

As Snf1 is phosphorylated in cells with dysfunctional mitochondria, we hypothesized that OXPHOS mutations might affect peroxisomal gene expression downstream of Snf1 activation, such as inactivation of the transcription inhibitors and/or the activation of the transcription activators. In addition, due to the lack of expression of some peroxisomal proteins in *ΔnugM* cells in glucose-limited conditions with prior induction by methanol, we surmised that the transcriptional activators, Prm1 and Mit1, induced by methanol should not be involved [73]. Nevertheless, we analyzed the role of Mit1 (Prm1 was excluded since it acts upstream of Mit1 in the same pathway [73]), which functions during methanol induction and the SNF1-regulated factor, Mxr1, functioning primarily during glucose depression.

We followed the expression of Pot1 and Aox1, setting aside the analysis of Pex11, whose regulation seems to mimic that of Pot1 and for which there is no antibody available. In addition, as gene expression is extremely complex, can be regulated at many levels and often includes the integration of many signals, to simplify our search for the mechanism underlying the OXPHOS deficiency and peroxisomal expression, we performed the following studies using either oleate or methanol induction, which include signals from glucose derepression, as well as induction by the new carbon source.

Contrary to our expectation (Fig. 4D) that inactivation of these transcription factors might mimic the OXPHOS deficiency, we found that none of the transcription activator deletions, *Δmxr1* or *Δmit1Δ*, shared the phenotype of the *ΔnugM* strain (Fig. 5). During oleate induction, both *Δmxr1* and *Δmit1* strains showed only a minor defect in Pot1 expression in comparison to WT cells, which was distinct from the behavior of the *ΔnugM* strain (Fig. 5A). However, during growth in methanol, both transcription activators were essential for Aox1 expression, as described previously [73, 74, 80] (Fig. 5B), confirming again the synergy of signals from the glucose derepression (SNF1 pathway) and from the methanol induction (Prm1 and Mit1) needed to activate the *AOX1* promoter.

**Figure 5:**
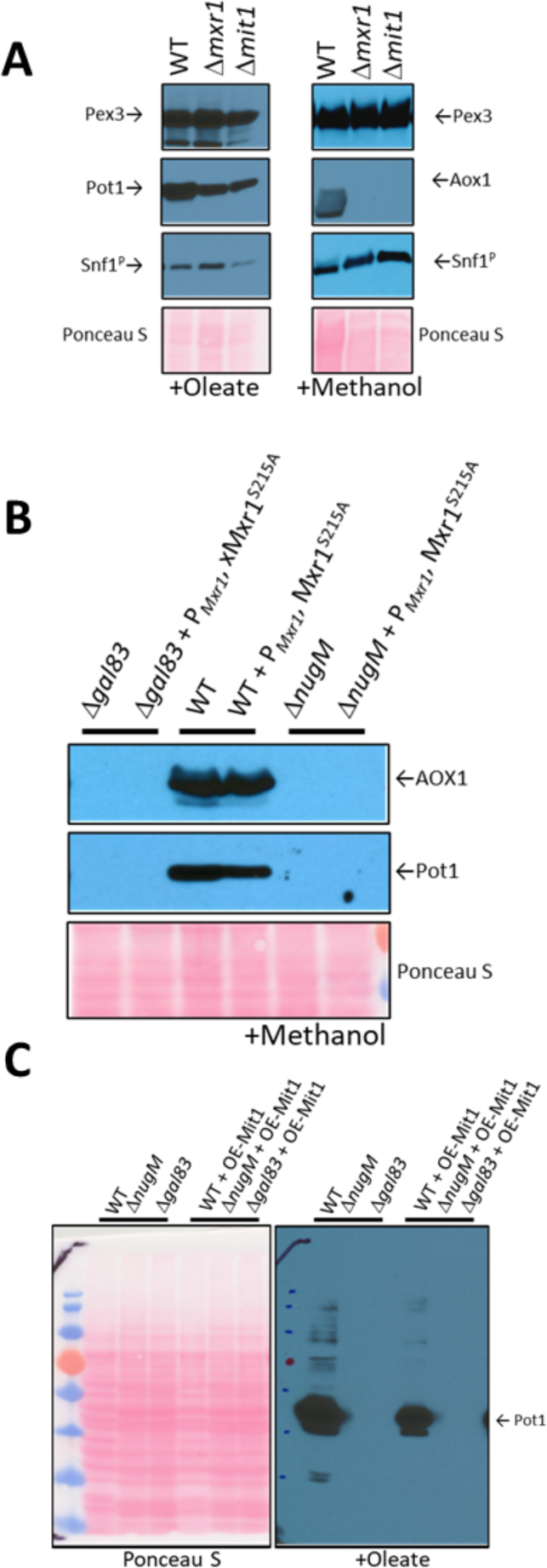
Analysis of transcriptional activators, Mxr1 and Mit1, of peroxisomal proteins. (A) Western blots of Pex3, phospho-Snf1, and Pot1 or Aox1 in WT, *Δmxr1* and *Δmit1* mutant cells. (B) Western blots of Pot1 and Aox1 in WT, *Δgal83* and *ΔnugM* mutant cells, either expressing or not expressing the active form of Mxr1 (Mxr1^S215A^). (C) Western blot of Pot1 in WT, *Δgal83* and *ΔnugM* mutant cells, without or with overexpression of Mit1 (OE-Mit1). Ponceau S staining was used as a loading control.

The role of these transcription activators was studied further by direct activation of the factors, bypassing their upstream regulatory steps. The activation mechanism of both transcription activators has been described. Briefly, during glucose derepression, the constitutively-expressed Mxr1 is activated by dephosphorylation at Ser 215 [80] and during methanol induction, Prm1 transmits the methanol signal to Mit1 by binding to the *MIT1* promoter, thus increasing expression of Mit1 [73].

To accomplish constitutive activation of the TFs, we mutated the Mxr1 phosphorylation site from Ser 215 to Ala (Mxr1^S215A^), PKA was inhibited (PKA mutants are susceptible to 1NM-PP1 inhibition), which is the putative kinase for the inhibitory phosphorylation of the *S. cerevisiae* homolog of Mxr1 (Adr1) and Mit1 was overexpressed (OE-Mit1) from the strong, constitutive *GAPDH* promoter. We expected that if any of these transcription activators was not activated in the OXPHOS mutants, their constitutive activation should completely or partially rescue the peroxisomal protein expression defect seen in the OXPHOS mutants.

The plasmids expressing OE-Mit1, PKA mutants and Mxr1^S215A^ were transformed into WT, *Δgal83* and *ΔnugM* strains, respectively, and analyzed for the expression of Pot1 and Aox1, after induction by oleate or methanol (Fig. 5B, 5C and Fig. S4). OE-Mit1 reduced the cell fitness independent of the background strain and cells grew slower in all carbon sources tested. Constitutive Mxr1 activation (Mxr1^S215A^) did not affect the fitness of cells, but like OE-Mit1, did not rescue the OXPHOS or SNF1 deficiencies (Fig. 5B). Independent of the growth defect, the overexpression of Mit1 did not rescue Aox1 or Pot1 expression in *ΔnugM* or *Δgal83* cells, during methanol (not shown) or oleate induction (Fig. 5C). Similar results were obtained when we inhibited the putative Mxr1 kinase, PKA (Fig. S4).

Aox1 is substantially induced when cells are cultured in methanol and constitutes 30% of total soluble proteins in the yeast cell [81]. While strongly induced by methanol, the *AOX1* promoter is strictly repressed by other carbon sources such as glucose, glycerol and ethanol [82]. Recently, the *P. pastoris* MAP kinase, Hog1, was implicated in Aox1 repression under glycerol conditions [77] and in *S. cerevisiae*, Snf1 negatively regulates the activity of Hog1 [83]. However, deletion of the *HOG1* gene in *ΔnugM* cells did not rescue the expression of Pot1 and Aox1 (Fig. S4).

### Rescue of the OXPHOS defect by inactivation of transcriptional repressors of genes encoding peroxisomal proteins

In contrast to the transcriptional activator ScAdr1, the regulation of the transcription inhibitors, Mig1, Mig2 and Nrg1 by the SNF1 complex in *S. cerevisiae* has been clearly established, with Snf1 kinase directly phosphorylating Mig1 [41] and Mig2 [44], or directly interacting with Nrg1 [84]. In *S. cerevisiae*, the direct phosphorylation of Mig1 and Mig2 by the active SNF1 complex causes their release from the promoter of glucose-repressed genes, followed by their export to the cytosol.

In *P. pastoris*, the triple deletion strain (*Δmig1 Δmig2 Δnrg1*) strongly derepressed Aox1 under glycerol-repression conditions, but not during glucose-derepression, even when transcription activators (Mit1 and Prm1) were artificially activated [85]. This makes sense because deletion of *MIG1* and *MIG2* significantly upregulated the binding of Mit1 at the *AOX1* promoter [71].

We used the already available *Δmig1 Δmig2 Δnrg1* strain, integrating the peroxisomal marker Pex3-GFP, and we then deleted the *NUGM* gene, and *GAL83* gene as a control. Most of the peroxisome proliferation defects we observed in the OXPHOS and *gal83* mutants, such as peroxisome proliferation, as well as Pot1 and Aox1 expression, were rescued when cells were induced in either oleate or methanol (Fig. 6A, B). Both quadruple mutants (*Δmig1 Δmig2 Δnrg1 ΔnugM* and *Δmig1 Δmig2 Δnrg1 Δgal83*) behaved similarly in both media. Additionally, peroxisome proliferation, assessed by fluorescence microscopy of Pex3-GFP, was identical to that in WT and the triple mutant (*Δmig1 Δmig2 Δnrg1*) strains (Fig. 6C and 6D). We observed a reduction in Aox1 expression during oleate induction (glucose derepression) in the *Δmig1 Δmig2 Δnrg1 ΔnugM* strain, in comparison with the WT and the triple mutant strains, but this difference was also observed in *Δmig1 Δmig2 Δnrg1 Δgal83* strain (Fig. 6A), and was therefore considered insignificant. Thus, most of the peroxisome-associated defects caused the loss of mitochondrial ATP production can be rescued by inactivation of the three transcriptional repressors, closing the mechanistic loop involved in mitochondria-peroxisome interplay.

**Figure 6:**
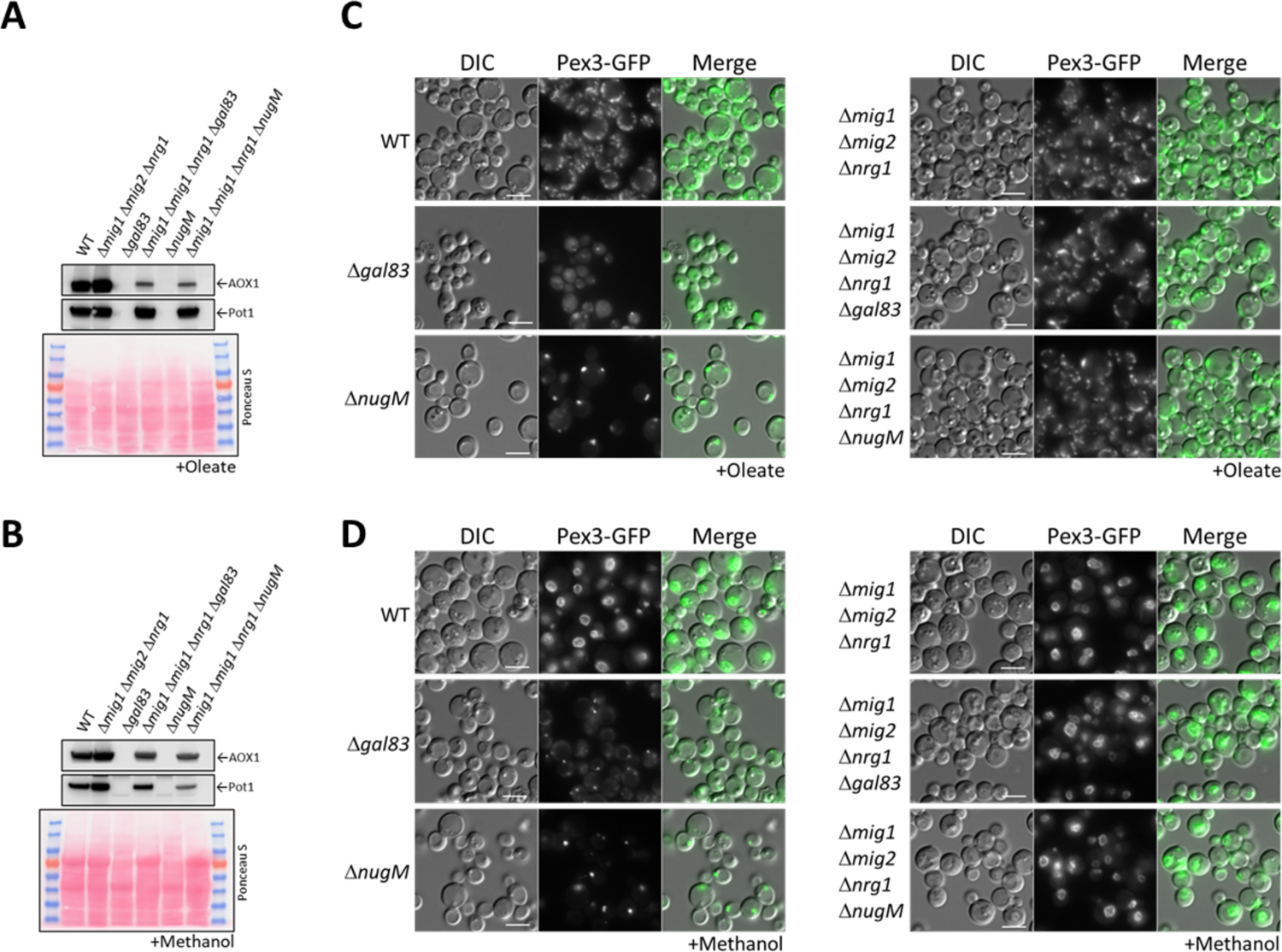
Deletion of transcriptional repressors regulated by SNF1 complex signaling rescues **Δ*nugM* and Δ*gal83* mutant cells.** (A) and (B) Western blot of Aox1 and Pot1 in WT, *ΔnugM* and *Δgal83* mutant cells, with and without deletions of genes encoding the transcriptional repressors regulated by SNF1 signaling (*MIG1*, *MIG2* and *NRG1*). Ponceau S staining was used as a loading control. (C) and (D) Fluorescence microscopy of WT, *Δgal83* and *ΔnugM* mutant cells expressing Pex3-GFP driven by the *PEX3* promoter, with and without deletions of genes encoding the transcriptional repressors regulated by SNF1 signaling (*MIG1*, *MIG2* and *NRG1*). Cells were grown in oleate and methanol for 16 h, respectively. Bars: 5 μm.

### Gal83 nuclear shuttling during peroxisome proliferation is inhibited in OXPHOS mutants

Transcriptional repressors and activators shuttle between the cytoplasm and the nucleus in a glucose-dependent manner. However, the mechanism regulating the traffic remains largely unknown. Similarly, the SNF1 complex moves to the nucleus to inactivate and to activate these repressors and activators, respectively. There is no doubt that the SNF1 complex needs to be localized in the nucleus to inactivate the repressors that occlude the promoters of glucose-repressed genes, but there is little to no evidence for Mxr1/Adr1 activation in the nucleus. Instead, if Mxr1/Adr1 indeed can be activated within the cytosol, a deficient nuclear enrichment of SNF1 in the OXPHOS mutant during glucose derepression could explain the rescue of peroxisome biogenesis defects in the *ΔnugM* mutant by deletion of the repressors. In *S. cerevisiae*, SNF1 complex localization is regulated by the *β*-subunits (Gal83, Sip1 and Sip2), with Gal83 being the subunit responsible for the nuclear localization of the complex (referred to hereafter as SNF1-Gal83).

We tested for effects of the of *ΔnugM* mutant on Gal83 nuclear localization by using a Gal83-GFP fusion expressed from the native *GAL83* promoter. WT and mutant cells were grown on abundant glucose, and the nuclear enrichment of Gal83-GFP was stimulated by shifting the cells to oleate as the carbon source for 30 min. As expected, Gal83-GFP was excluded from the nuclei of glucose-grown WT and *Δsak1* cells (Fig. 7 and Fig. S5), and similarly excluded from the nuclei of the *ΔnugM* mutant (Fig. 7). Upon the shift to oleate, Gal83-GFP was enriched in the nucleus of the WT and excluded from the nucleus of the *Δsak1* mutant, as anticipated (Fig. 7). However, no nuclear enrichment was observed in *ΔnugM* cells (Fig. 7 and Fig. S5). Together with the Snf1 activation assay (Fig. 4D), these results strongly suggest that the OXPHOS mutants are impaired in the nuclear localization of the SNF1-Gal83 complex by a mechanism that is independent of the catalytic activation of Snf1 kinase.

**Figure 7:**
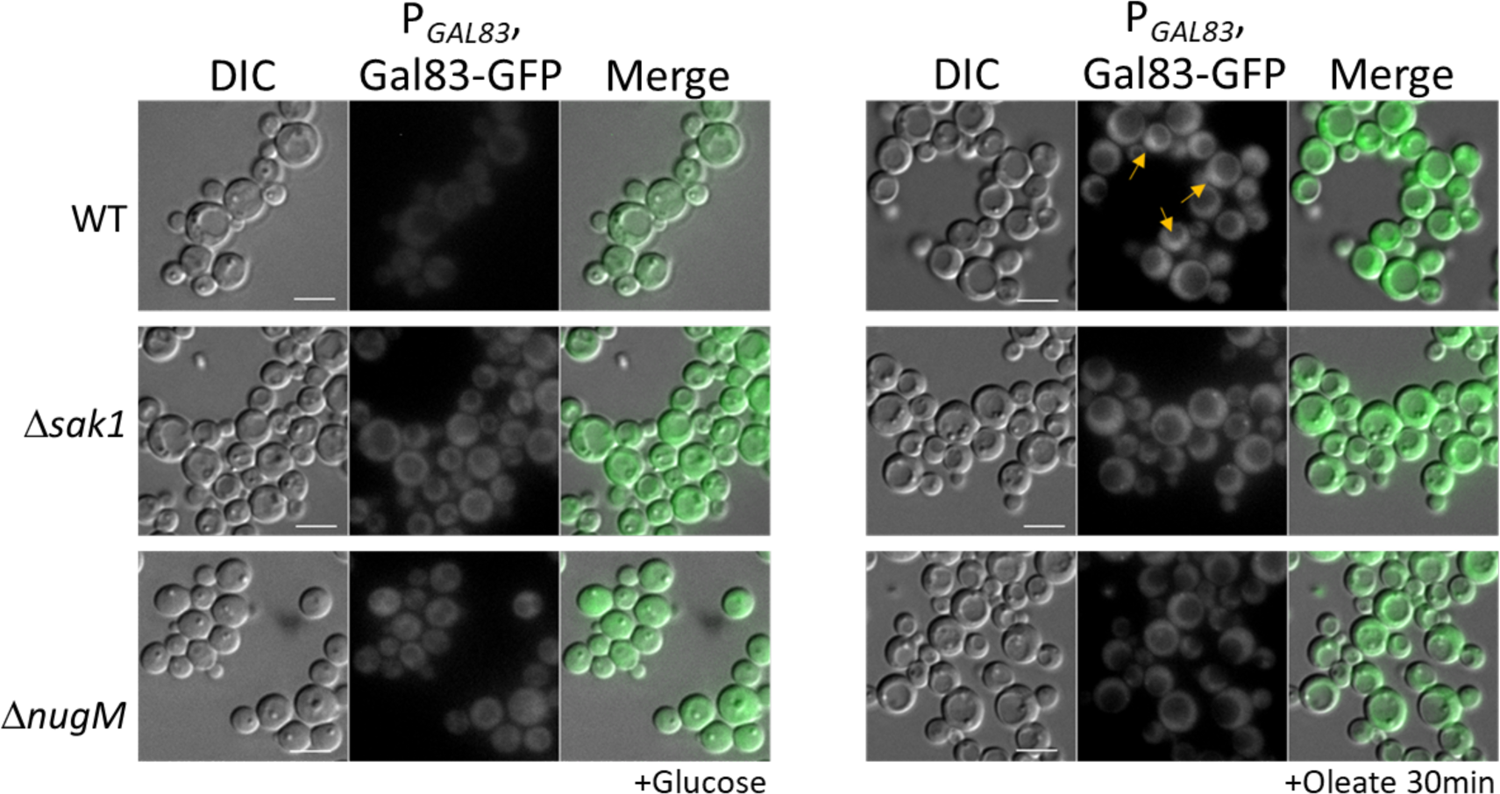
Gal83 nuclear localization during oleate adaptation is inhibited in Δ*nugM* and **Δ*sak1* mutant cells.** Fluorescence microscopy of WT, *ΔnugM* and *Δgal83* mutant cells expressing Gal83-GFP driven by the *GAL83* promoter. Bars: 5 μm.

### Peroxisome proliferation in Δ*pex14* cells can be induced by growth in lactate medium

The aberrant regulation of the SNF1 signaling pathway in cells with dysfunctional mitochondria during glucose derepression, and the lack of a normal peroxisome proliferation in NADH shuttling mutants during growth in oleate and methanol suggest the presence of a feedback loop between both organelles which goes beyond the initial activation of the SNF1 signaling pathway by glucose derepression.

We tested the feedback loop hypothesis using a strain lacking Pex14, a key member of the peroxisome docking complex and essential for import of peroxisomal matrix proteins [86]. Although normal peroxisomes are absent in cells lacking Pex14, abnormal vesicles containing peroxisomal membrane proteins (PMPs) but lacking matrix proteins, are observed as peroxisomal remnants. These remnants can be visualized with another PMP, Pex3-GFP, expressed from its own promoter, and when we compared *Δpex14* mutant to WT cells after cultivation in oleate medium, we observed a significantly lower number of peroxisomal structures (Fig. 8). We investigated if an inactive OXPHOS (due to the lack of NADH production by dysfunctional peroxisomes) is the reason for the low number of remnants. We noticed after 4 h of cultivation in L-lactate (lactate) medium (non-fermentable carbon source), that WT cells showed higher levels of Pex11, Aox1 and Pot1 proteins compared to glucose depletion or to cultivation in methanol or oleate (Fig. 4D), which could be a consequence of the lactate metabolism by the mitochondria. Therefore, we used the lactate medium to overcome the low levels of NADH in *Δpex14* cells, and as we hypothesized, the numbers of peroxisome structures increased in *Δpex14* cells and were comparable to those in WT cells cultivated in lactate medium. Additionally, this result suggests that peroxisome proliferation cannot be dependent solely from a peroxisomal metabolite because *Δpex14* cells would be unable to produce these.

**Figure 8:**
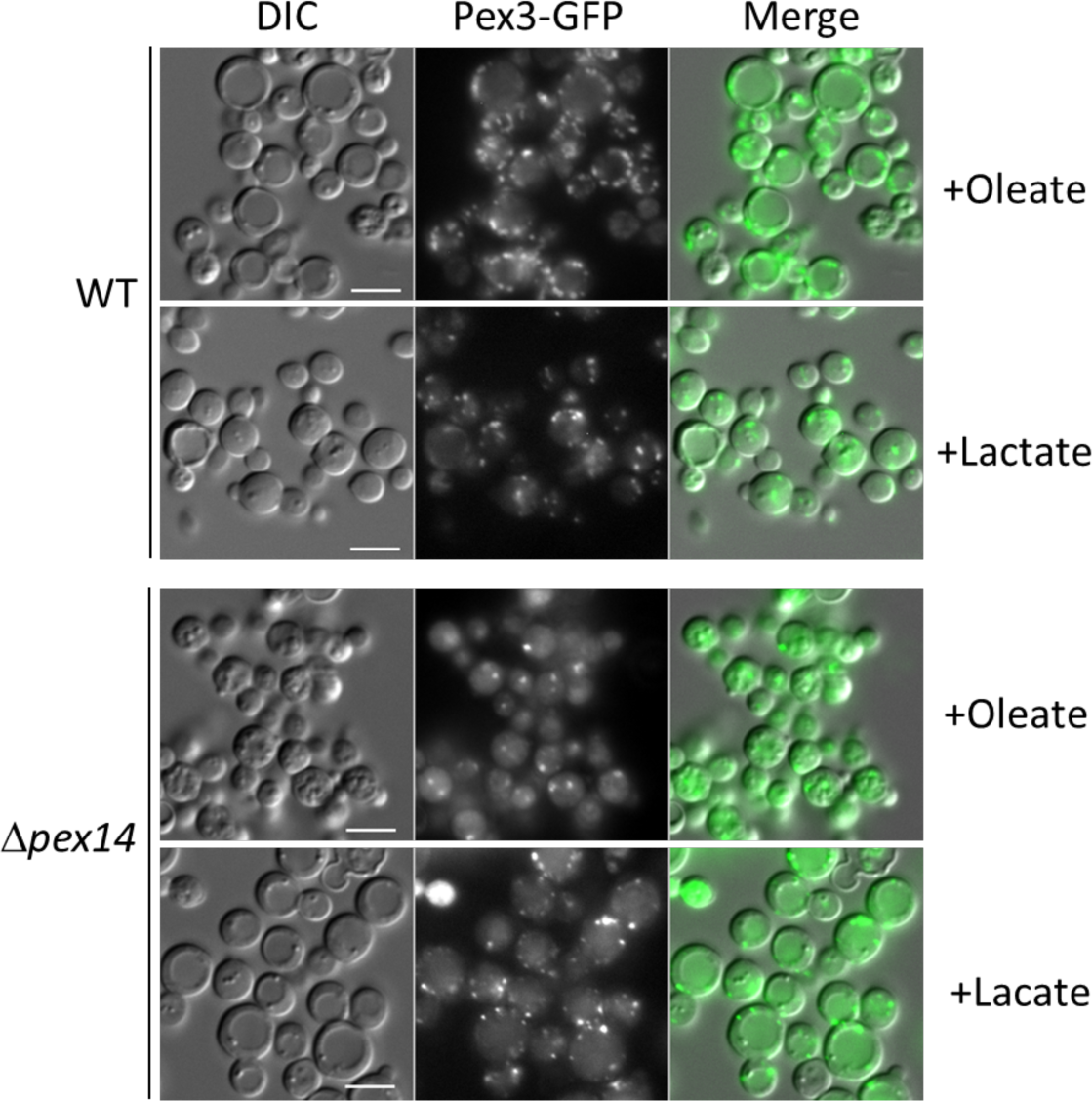
Cultivation of Δ*pex14* mutant cells in the respiratory medium, lactate, induces peroxisome proliferation. Fluorescence microscopy of WT, *Δpex14* mutant cells expressing Pex3-GFP driven by the *PEX3* promoter in different media for 8 h. Bars: 5 μm.

## Discussion

Organelles are often viewed as individual entities with defined composition and organization that endow them with specialized functions. However, it is now clear that intracellular membrane compartments engage in extensive communication, either indirectly, or directly through membrane contacts [52, 87–90]. Mitochondrial dysfunction impacts several other organelles and their biogenesis, including peroxisomes [91], but the underlying mechanisms have not been elucidated. We have uncovered here the mechanisms involved in one mode of interorganelle communication and interplay in *P. pastoris* where the mitochondria and peroxisomes sense the metabolic status of the cell, influence each other’s metabolism, in concert with cytosolic and nuclear involvement, and regulate peroxisome proliferation and division as needed.

### Interorganellar communication and interplay between peroxisomes and mitochondria

In the mitochondrial OXPHOS mutants (*ΔnugM, Δndufa9 and Δcyt1),* and upon mitochondrial uncoupling using DNP, peroxisome proliferation, division and biogenesis of several peroxisome-associated proteins, but not peroxisomal matrix protein import (Fig. 2), are affected in cells grown in either methanol or oleate, illustrating the mitochondrial involvement in multiple peroxisomal processes. We extended earlier reports that intermediates of peroxisome metabolism, or the absence of peroxisomal enzymes, regulate the maturation and fission of the organelle [56, 57] by several additional observations. The lack of peroxisomal thiolase (Pot1) affected peroxisome proliferation and division exclusively in oleate. Similarly, the absence of Aox1 and Aox2, involved in the methanol utilization pathway, affected peroxisome proliferation and division only in cells grown in methanol medium (Fig. S1A). Finally, the lack of Pex5, responsible for the import of PTS1-containing enzymes (including some involved in the *β*-oxidation and methanol metabolism pathways), impaired peroxisome proliferation and division in both media (Fig. S1B). Additionally, blocking the entry of NADH into mitochondria using double deletion strains, *ΔmdhA Δgpd1* and *ΔmdhB Δgpd1* cells, yielded a phenotype like that seen in the OXPHOS mutants, wherein most of the cells contained only a single, import-competent peroxisome (Fig. 1C). These results are consistent with the role of NADH-shuttling proteins, responsible for shuttling NADH between both peroxisomes and mitochondria via the cytosol, in the interorganellar interplay between these organelles, and also points to a role for the cytosol.

### Feedback loop between peroxisomes and mitochondria senses cellular metabolic status

Our results in *P. pastoris* during glucose derepression indicate that peroxisome proliferation, division and biogenesis rely on a functional OXPHOS. Moreover, as methylotrophic yeast rely exclusively on peroxisome metabolism to feed into the mitochondrial OXPHOS for energy production during growth in FA or methanol [92, 93], this feedback loop between the two organelles might contribute to sensing the cell’s metabolic status during cultivation in these carbon sources (Fig. 9). We obtained proof of this feedback loop when we induced the peroxisome proliferation in *Δpex14* cells, which cannot import peroxisomal matrix proteins and produce NADH, by cultivation in lactate, a non-fermentable carbon source (Fig. 8). In yeast, lactate can only be metabolized by the mitochondria, and any externally-added lactate will enter mitochondria via a putative lactate/proton symporter and will then be oxidized to pyruvate with a reduction of cytochrome C in an energy competent manner [94]. In *Δpex14* cells, cultivation in lactate activated OXPHOS by mitochondrial metabolism and overrode the need for NADH produced by peroxisomal metabolism, boosting the peroxisome proliferation (Fig. 8). This also shows clearly that the molecule triggering peroxisome proliferation does not emanate from within peroxisomes, which are dysfunctional in *Δpex14* cells.

**Figure 9:**
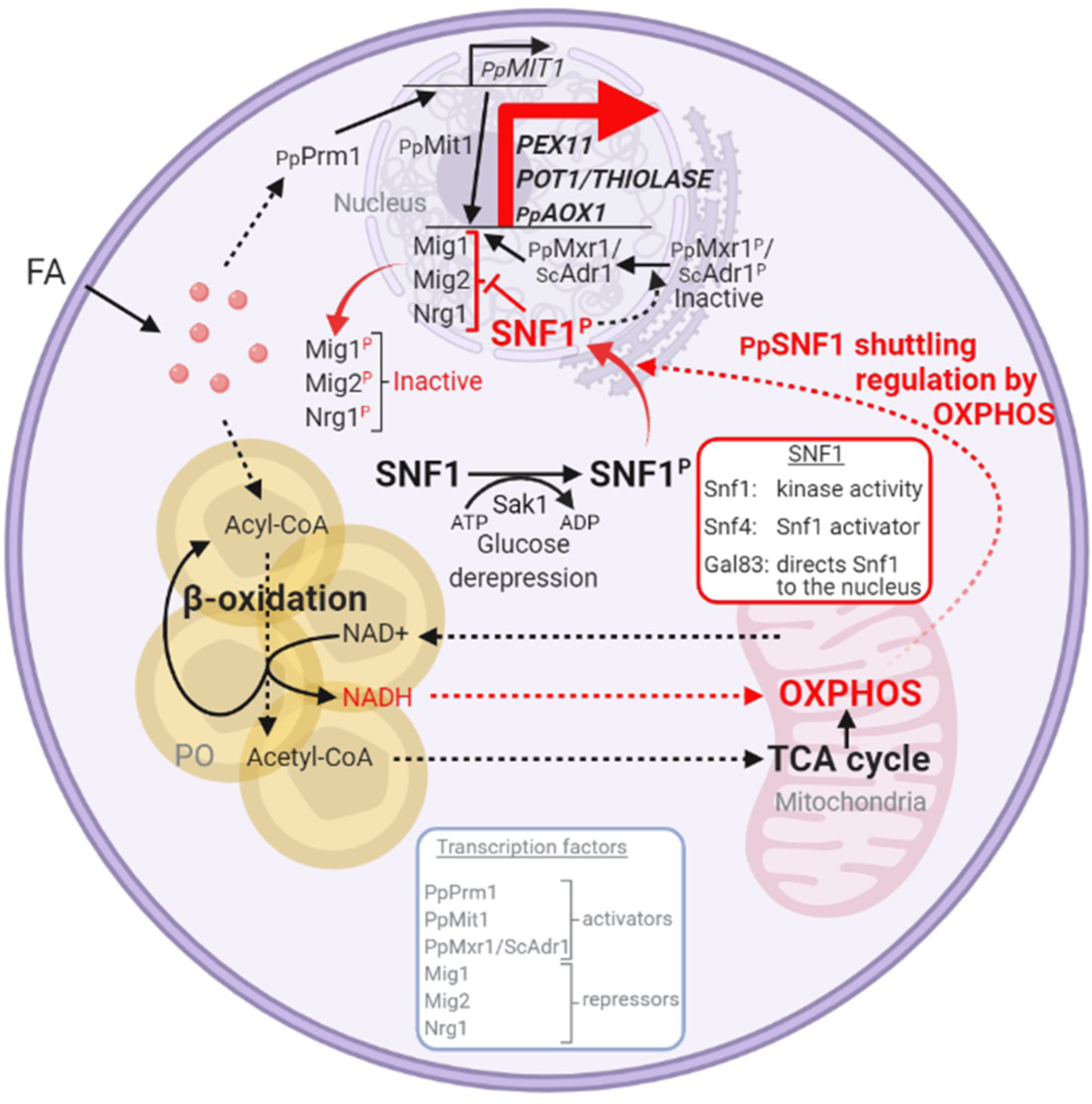
Interorganellar communication and signaling pathways in peroxisome proliferation, division and matrix protein biogenesis. Feedback loop between peroxisome and mitochondria is shown in red. FA, fatty acids. PO, peroxisome. P: phosphorylation. See text for details.

### The OXPHOS effect on peroxisome phenotypes in mediated by the absence of SNF1-Gal83 translocation to the nucleus

Despite the pleiotropic effects of the mitochondrial OXPHOS mutants on multiple peroxisome properties like proliferation, division and biogenesis of peroxisome-associated proteins, it was interesting that most of these phenotypes were not caused not by the lack of NADH *per se*, but rather the reduced ATP production in the OXPHOS mutants is the key, because the same phenotype was obtained using the uncoupler, DNP (Fig. 3).

More specifically, the effect of ATP is likely mediated through the SNF1 complex that is known to respond to altered cellular AMP/ATP ratios. Our data show that the phenotype of the OXPHOS mutants was due to the lack of nuclear translocation of the SNF1-Gal83 complex (Fig. 7) and a defect in the transcriptional induction of peroxisome division (Pex11) and peroxisomal matrix proteins (e.g. Pot1 and Aox1) in the OXPHOS mutants grown in methanol or oleate (Fig. 4D). This reiterates a nuclear involvement in the activation of peroxisome proliferation.

We present several lines of evidence showing that the newly-discovered role played by OXPHOS in nuclear enrichment of the SNF1-Gal83 complex is separable from its role in catalytic activation of the kinase, and it rather reflects a role in regulating the nuclear relocalization of Gal83 itself. First, the Δ*nugM* mutation alone does not affect Thr 210 phosphorylation and activation of Snf1 kinase (Fig. 4D). Second, Gal83 fails to enrich in the nucleus of the Δ*nugM* mutant, under conditions where this relocation is evident in WT cells (Fig. 7). The pathway leading to Gal83 import and the proteins involved in this process are not known. Gal83 may undergo a posttranslational modification upon glucose derepression that could promote a change in cellular localization. In *S. cerevisiae*, Gal83 phosphorylation sites have been identified in high-throughput studies [95], but the effect of such modifications is not known. Some evidence suggests a role for the mitochondrial voltage-dependent anion channel (VDAC) protein, Por1, promoting Gal83 nuclear localization by a mechanism that is distinct from Snf1 activation during growth in glycerol/ethanol [96], and for the nuclear export receptor, Crm1, in the nuclear exclusion of Gal83 during growth on abundant glucose [97]. *P. pastoris* OXPHOS mutants share phenotypes of *ΔScpor1* mutant, such as normal Snf1 activation by glucose derepression and the absence of induction of genes regulated by SNF1 signaling [96]. However, the deletion of the only *POR1* homolog, or overexpression of Por1 in *P. pastoris,* did not produce the same phenotypes observed in the OXPHOS mutants and Gal83 localization was unaffected by Por1. Corroborating the role of Gal83 was also the finding the *Δgal83* cells phenocopied the OXPHOS mutants (Fig. 4C).

In view of the fact that AMPK enhances PGC-1α expression and activity, the role of master transcriptional co-regulators of peroxisomal and mitochondrial proteins is not unexpected [98]. In *S. cerevisiae*, and mammals, the Snf1/AMPK signaling pathway co-induces several mitochondrial and peroxisomal genes [67, 99].

### The downstream targets of the SNF1-Gal83 signaling explain the peroxisome-related phenotypes in the OXPHOS mutants

Among the proteins whose expression is affected transcriptionally in the OXPHOS mutants, is Pex11. The Pex11 family of proteins is conserved in yeasts, plants and mammals, and orchestrates peroxisome division [19, 20, 23, 100, 101]. Its absence blocks peroxisome division in yeast, but importantly, not its *de novo* proliferation [18], which is seen quite clearly in *P. pastoris* where *Δpex11* mutants grown in oleate have only 1-2 large peroxisomes, in comparison to WT cells that have about 7 peroxisomes/cell [21]. However, in methanol there is indeed peroxisome proliferation and cells have multiple peroxisomes in cells lacking either Pex11 or its downstream peroxisome division component, Fis1 (unpublished data cited in [21]). Additionally, the *P. pastoris* Δ*pex11* mutant does not phenocopy the OXPHOS mutant (Fig. S6). In both *S. cerevisiae* and *P. pastoris*, Pex11 is induced significantly [29] upon switch from YPD to oleate [21]. In *P. pastoris*, Pex11 is fully repressed upon growth of cells in YPD, whereas in *S. cerevisiae,* Pex11 is expressed at a low level even in YPD [21]. The absence of induction of Pex11 in oleate-grown cells (Fig. 4A, D) is likely the underlying cause for the lack of peroxisome division in the OXPHOS mutants, but would not explain the additional absence of peroxisome proliferation.

The biogenesis defects we observed for peroxisome-associated proteins in the OXPHOS mutants are likely caused by the lack of induction of peroxisomal matrix proteins, such as Pot1 and Aox1 (Fig. 4C-E), and their corresponding RNAs (Fig, 4D). Further corroboration of this conclusion comes from our data that the absence of several peroxisomal matrix proteins (Pot1, Aox1 and Aox2) by deletion of their corresponding genes, or by prevention of their import into peroxisomes (*PEX5* gene deletion) also affects peroxisome proliferation, as seen for the OXPHOS mutants (Fig. S1A, S1B).

Finally, the rescue of the phenotype of the OXPHOS mutant by deletion of the genes encoding the repressors *MIG1*, *MIG2* and *NRG1*, extends not only to the induction of *PEX11*, *POT1* and *AOX1* genes (Fig. 6A, B), but also to the reversion of the defect in peroxisome proliferation (Fig. 6C, 6D), making it very likely that the missing factor required for peroxisome proliferation in the OXPHOS mutants must be a direct or indirect target of these transcriptional repressors, but as argued earlier, based on the peroxisome proliferation seen in *Δpex14* cells in lactate, this is unlikely to involve a peroxisomal metabolite. In contrast, peroxisome division can be influenced by peroxisomal metabolites [102]. Identification of this proliferation factor remains an important future priority.

In support of our conclusion that most of the peroxisome-related phenotypes could be explained at the transcriptional level via the action of the SNF1-Gal83 signaling pathway, we were able to rescue the peroxisome proliferation, peroxisomal protein biogenesis and division defects in the OXPHOS and *Δgal83* mutants, by the simultaneous deletion of three genes encoding the transcriptional repressors Mig1, Mig2 and Nrg1 (Fig. 6), which are known to repress genes required for peroxisome biogenesis and function [71, 76]. Deletions of the genes for the transcriptional activators, Mit1 and Mxr1, known to activate peroxisome-related genes [73, 74], individually, did not yield the same phenotype as that of the OXPHOS mutants (Fig. 5A), suggesting that these are not directly impaired in the OXPHOS mutants. Additionally, neither overexpression of Mit1, nor artificial activation of Mxr1, rescued the similar phenotypes of the *ΔnugM* and *Δgal83* mutants (Fig. 5B, C). Therefore, most of the peroxisome-related phenotypes of the OXPHOS mutants can be explained by the lack of relief of repression of several genes by Mig1, Mig2 and Nrg1, thereby providing a molecular explanation for how the ATP generated by the mitochondria impact peroxisome proliferation, division and biogenesis.

### Working model for interorganellar control of peroxisome dynamics

We present a working model illustrating the interorganellar transactions, communications and signaling that must occur in *P. pastoris* between peroxisomes, cytosol, mitochondria and the nucleus (Fig. 9) for peroxisome proliferation, division and biogenesis of peroxisome-associated proteins. FA uptake and its β-oxidation produce NADH equivalents and acetyl-CoA. However, the peroxisome membrane is impermeable to large hydrophilic solutes, including NAD^+^, NADH, NADP^+^ and NADPH, as well as ATP, and either acylated or unacylated coenzyme A (CoA) [2]. Consequently, NADH-shuttling proteins, working together in the peroxisomes, cytosol and mitochondria, allow the delivery of NADH to mitochondria to feed OXPHOS [64]. Acetyl-CoA produced in peroxisomes is delivered to mitochondria via acyl-carnitine produced in peroxisomes, and mitochondria use the TCA cycle and OXPHOS system for full oxidation to CO_2_ and H_2_O [2].

The SNF1/AMP-activated protein kinase (AMPK) complex, which is sensitive to the cellular AMP:ATP ratio, maintains the balance between ATP production and consumption in all eukaryotic cells [68, 69, 79]. Glucose deprivation, which reduces ATP production, activates Snf1 (by phosphorylation) via the action of the Sak1 kinase, and in *P. pastoris*, the nuclear translocation of the SNF1-Gal83 complex requires OXPHOS and ATP production, as shown here. Peroxisome-associated proteins, such as the division protein, Pex11, and the matrix proteins, Pot1 and Aox1, are regulated negatively by transcriptional repressors, that compete with transcriptional activators, such as Mit1 and Mxr1 (equivalent to ScAdr1). Snf1 activation in the cytosol and SNF1-Gal83 entry into the nucleus removes, by phosphorylation of the appropriate proteins, the repression of expression of the peroxisome-associated proteins, while also activating the transcriptional activators. This simultaneous action of SNF1-Gal83 turns on the biogenesis of peroxisome-associated proteins, peroxisome proliferation, as well as division. In the Δ*nugM*, Δ*ndufa9*, Δ*gal83*, Δ*pot1*, Δ*aox1* Δ*aox2* and Δ*pex5* mutants of *P. pastoris*, or in the presence of the mitochondrial uncoupler, DNP, peroxisome proliferation, division and the biogenesis of certain peroxisome-associated proteins in compromised.

In conclusion, since ATP production by mitochondrial OXPHOS is impaired in many human diseases involving over 150 genes [103], including Parkinson’s disease and schizophrenia [104–106], as well as during aging [107] and neurodegeneration [108], the results presented here provide a clear mechanistic link to peroxisomal dysfunction in human mitochondrial disorders. Since SNF1/AMPK activation in both yeast and mammalian cells upregulates glycolysis, gluconeogenesis and mitochondrial biogenesis, while repressing ATP consumption [103, 109], it makes sense that this would also enhance peroxisome proliferation because peroxisomes house several glyoxylate pathway enzymes that are necessary for gluconeogenesis [110]. Likewise, since peroxisomal metabolites (like NADH and products of the *β*-oxidation of very long chain FA) influence mitochondrial ATP production [111], these studies also explain how diseases affecting peroxisome biogenesis can impair mitochondria. Further explorations of this yeast model can provide important insights regarding interorganellar communication, interplay and dynamics, while also shedding light on human disease.

## Acknowledgements

This research was funded by the NIH grant (RO1 DK41737) to SS, who holds a Tata Chancellor’s Endowed Professorship in Molecular Biology. We thank Dr. Barry Bochner and In Iok Kong from Biolog Inc., Hayward, CA for advice and use of their Biolog machine.

## References

1. Lazarow PB (2003) Peroxisome biogenesis: advances and conundrums. Curr Opin Cell Biol 15: 489–97

2. Wanders RJA, Vaz FM, Waterham HR, Ferdinandusse S (2020) Fatty Acid Oxidation in Peroxisomes: Enzymology, Metabolic Crosstalk with Other Organelles and Peroxisomal Disorders. Adv Exp Med Biol 1299: 55–70

3. Sutterlin C, Hsu P, Mallabiabarrena A, Malhotra V (2002) Fragmentation and dispersal of the pericentriolar Golgi complex is required for entry into mitosis in mammalian cells. Cell 109: 359–69

4. Osteryoung KW, Nunnari J (2003) The division of endosymbiotic organelles. Science 302: 1698–704

5. Lingard MJ, Gidda SK, Bingham S, Rothstein SJ, Mullen RT, Trelease RN (2008) Arabidopsis PEROXIN11c-e, FISSION1b, and DYNAMIN-RELATED PROTEIN3A cooperate in cell cycle-associated replication of peroxisomes. Plant Cell 20: 1567-85

6. Fagarasanu A, Rachubinski RA (2007) Orchestrating organelle inheritance in Saccharomyces cerevisiae. Curr Opin Microbiol 10: 528–38

7. Hettema EH, Motley AM (2009) How peroxisomes multiply. J Cell Sci 122: 2331–6

8. Guo T, Kit YY, Nicaud JM, Le Dall MT, Sears SK, Vali H, Chan H, Rachubinski RA, Titorenko VI (2003) Peroxisome division in the yeast Yarrowia lipolytica is regulated by a signal from inside the peroxisome. J Cell Biol 162: 1255–66

9. Veenhuis M, Kiel JA, Van Der Klei IJ (2003) Peroxisome assembly in yeast. Microsc Res Tech 61: 139–50

10. Farre JC, Subramani S (2016) Mechanistic insights into selective autophagy pathways: lessons from yeast. Nat Rev Mol Cell Biol 17: 537–52

11. Chang CC, South S, Warren D, Jones J, Moser AB, Moser HW, Gould SJ (1999) Metabolic control of peroxisome abundance. J Cell Sci 112 (Pt 10): 1579–90

12. Poll-The BT, Roels F, Ogier H, Scotto J, Vamecq J, Schutgens RB, Wanders RJ, van Roermund CW, van Wijland MJ, Schram AW, et al. (1988) A new peroxisomal disorder with enlarged peroxisomes and a specific deficiency of acyl-CoA oxidase (pseudo-neonatal adrenoleukodystrophy). Am J Hum Genet 42: 422–34

13. Smith JJ, Brown TW, Eitzen GA, Rachubinski RA (2000) Regulation of peroxisome size and number by fatty acid beta-oxidation in the yeast yarrowia lipolytica. J Biol Chem 275: 20168–78

14. Yan M, Rayapuram N, Subramani S (2005) The control of peroxisome number and size during division and proliferation. Curr Opin Cell Biol 17: 376–83

15. Schrader M (2006) Shared components of mitochondrial and peroxisomal division. Biochim Biophys Acta 1763: 531–41

16. Kuravi K, Nagotu S, Krikken AM, Sjollema K, Deckers M, Erdmann R, Veenhuis M, van der Klei IJ (2006) Dynamin-related proteins Vps1p and Dnm1p control peroxisome abundance in Saccharomyces cerevisiae. J Cell Sci 119: 3994–4001

17. Motley AM, Ward GP, Hettema EH (2008) Dnm1p-dependent peroxisome fission requires Caf4p, Mdv1p and Fis1p. J Cell Sci 121: 1633-40

18. Huber A, Koch J, Kragler F, Brocard C, Hartig A (2012) A subtle interplay between three Pex11 proteins shapes de novo formation and fission of peroxisomes. Traffic 13: 157–67

19. Koch J, Brocard C (2012) PEX11 proteins attract Mff and human Fis1 to coordinate peroxisomal fission. J Cell Sci 125: 3813–26

20. Koch J, Pranjic K, Huber A, Ellinger A, Hartig A, Kragler F, Brocard C (2010) PEX11 family members are membrane elongation factors that coordinate peroxisome proliferation and maintenance. J Cell Sci 123: 3389–400

21. Joshi S, Agrawal G, Subramani S (2012) Phosphorylation-dependent Pex11p and Fis1p interaction regulates peroxisome division. Mol Biol Cell 23: 1307–15

22. Knoblach B, Rachubinski RA (2010) Phosphorylation-dependent activation of peroxisome proliferator protein PEX11 controls peroxisome abundance. J Biol Chem 285: 6670–80

23. Aung K, Zhang X, Hu J (2010) Peroxisome division and proliferation in plants. Biochem Soc Trans 38: 817–22

24. Thoms S, Gartner J (2012) First PEX11beta patient extends spectrum of peroxisomal biogenesis disorder phenotypes. J Med Genet 49: 314–6

25. Wang J, Li L, Zhang Z, Qiu H, Li D, Fang Y, Jiang H, Chai RY, Mao X, Wang Y, et al. (2015) One of Three Pex11 Family Members Is Required for Peroxisomal Proliferation and Full Virulence of the Rice Blast Fungus Magnaporthe oryzae. PLoS One 10: e0134249

26. Weng H, Ji X, Naito Y, Endo K, Ma X, Takahashi R, Shen C, Hirokawa G, Fukushima Y, Iwai N (2013) Pex11alpha deficiency impairs peroxisome elongation and division and contributes to nonalcoholic fatty liver in mice. Am J Physiol Endocrinol Metab 304: E187–96

27. Gurvitz A, Rottensteiner H (2006) The biochemistry of oleate induction: transcriptional upregulation and peroxisome proliferation. Biochim Biophys Acta 1763: 1392–402

28. Scheckhuber CQ (2019) Characterization of survival and stress resistance in S. cerevisiae mutants affected in peroxisome inheritance and proliferation, Deltainp1 and Deltapex11. Folia Microbiol (Praha)

29. Karpichev IV, Small GM (1998) Global regulatory functions of Oaf1p and Pip2p (Oaf2p), transcription factors that regulate genes encoding peroxisomal proteins in Saccharomyces cerevisiae. Mol Cell Biol 18: 6560–70

30. Smith JJ, Marelli M, Christmas RH, Vizeacoumar FJ, Dilworth DJ, Ideker T, Galitski T, Dimitrov K, Rachubinski RA, Aitchison JD (2002) Transcriptome profiling to identify genes involved in peroxisome assembly and function. J Cell Biol 158: 259–71

31. Rottensteiner H, Kal AJ, Filipits M, Binder M, Hamilton B, Tabak HF, Ruis H (1996) Pip2p: a transcriptional regulator of peroxisome proliferation in the yeast Saccharomyces cerevisiae. EMBO J 15: 2924–34

32. Rottensteiner H, Wabnegger L, Erdmann R, Hamilton B, Ruis H, Hartig A, Gurvitz A (2003) Saccharomyces cerevisiae PIP2 mediating oleic acid induction and peroxisome proliferation is regulated by Adr1p and Pip2p-Oaf1p. J Biol Chem 278: 27605–11

33. Jiang R, Carlson M (1997) The Snf1 protein kinase and its activating subunit, Snf4, interact with distinct domains of the Sip1/Sip2/Gal83 component in the kinase complex. Mol Cell Biol 17: 2099-106

34. Yang X, Jiang R, Carlson M (1994) A family of proteins containing a conserved domain that mediates interaction with the yeast SNF1 protein kinase complex. EMBO J 13: 5878–86

35. Vincent O, Townley R, Kuchin S, Carlson M (2001) Subcellular localization of the Snf1 kinase is regulated by specific beta subunits and a novel glucose signaling mechanism. Genes Dev 15: 1104–14

36. Elbing K, McCartney RR, Schmidt MC (2006) Purification and characterization of the three Snf1-activating kinases of Saccharomyces cerevisiae. Biochem J 393: 797–805

37. Hedbacker K, Hong SP, Carlson M (2004) Pak1 protein kinase regulates activation and nuclear localization of Snf1-Gal83 protein kinase. Mol Cell Biol 24: 8255–63

38. Hong SP, Leiper FC, Woods A, Carling D, Carlson M (2003) Activation of yeast Snf1 and mammalian AMP-activated protein kinase by upstream kinases. Proc Natl Acad Sci U S A 100: 8839–43

39. Vallier LG, Carlson M (1994) Synergistic release from glucose repression by mig1 and ssn mutations in Saccharomyces cerevisiae. Genetics 137: 49–54

40. Serra-Cardona A, Petrezselyova S, Canadell D, Ramos J, Arino J (2014) Coregulated expression of the Na+/phosphate Pho89 transporter and Ena1 Na+-ATPase allows their functional coupling under high-pH stress. Mol Cell Biol 34: 4420–35

41. Treitel MA, Kuchin S, Carlson M (1998) Snf1 protein kinase regulates phosphorylation of the Mig1 repressor in Saccharomyces cerevisiae. Mol Cell Biol 18: 6273–80

42. Young ET, Dombek KM, Tachibana C, Ideker T (2003) Multiple pathways are co-regulated by the protein kinase Snf1 and the transcription factors Adr1 and Cat8. J Biol Chem 278: 26146–58

43. Ratnakumar S, Kacherovsky N, Arms E, Young ET (2009) Snf1 controls the activity of adr1 through dephosphorylation of Ser230. Genetics 182: 735–45

44. Chandrashekarappa DG, McCartney RR, Schmidt MC (2011) Subunit and domain requirements for adenylate-mediated protection of Snf1 kinase activation loop from dephosphorylation. J Biol Chem 286: 44532–41

45. Ludin K, Jiang R, Carlson M (1998) Glucose-regulated interaction of a regulatory subunit of protein phosphatase 1 with the Snf1 protein kinase in Saccharomyces cerevisiae. Proc Natl Acad Sci U S A 95: 6245–50

46. Mayer FV, Heath R, Underwood E, Sanders MJ, Carmena D, McCartney RR, Leiper FC, Xiao B, Jing C, Walker PA, et al. (2011) ADP regulates SNF1, the Saccharomyces cerevisiae homolog of AMP-activated protein kinase. Cell Metab 14: 707–14

47. Dakik P, Titorenko VI (2016) Communications between Mitochondria, the Nucleus, Vacuoles, Peroxisomes, the Endoplasmic Reticulum, the Plasma Membrane, Lipid Droplets, and the Cytosol during Yeast Chronological Aging. Front Genet 7: 177

48. Mattiazzi Usaj M, Brloznik M, Kaferle P, Zitnik M, Wolinski H, Leitner F, Kohlwein SD, Zupan B, Petrovic U (2015) Genome-Wide Localization Study of Yeast Pex11 Identifies Peroxisome-Mitochondria Interactions through the ERMES Complex. J Mol Biol 427: 2072–87

49. Shai N, Yifrach E, van Roermund CWT, Cohen N, Bibi C, L IJ, Cavellini L, Meurisse J, Schuster R, Zada L, et al. (2018) Systematic mapping of contact sites reveals tethers and a function for the peroxisome-mitochondria contact. Nat Commun 9: 1761

50. Baumgart E, Vanhorebeek I, Grabenbauer M, Borgers M, Declercq PE, Fahimi HD, Baes M (2001) Mitochondrial alterations caused by defective peroxisomal biogenesis in a mouse model for Zellweger syndrome (PEX5 knockout mouse). Am J Pathol 159: 1477–94

51. Salpietro V, Phadke R, Saggar A, Hargreaves IP, Yates R, Fokoloros C, Mankad K, Hertecant J, Ruggieri M, McCormick D, et al. (2015) Zellweger syndrome and secondary mitochondrial myopathy. Eur J Pediatr 174: 557–63

52. Fransen M, Lismont C, Walton P (2017) The Peroxisome-Mitochondria Connection: How and Why? Int J Mol Sci 18

53. Gould SJ, McCollum D, Spong AP, Heyman JA, Subramani S (1992) Development of the yeast Pichia pastoris as a model organism for a genetic and molecular analysis of peroxisome assembly. Yeast 8: 613–28

54. Cregg JM, Russell KA (1998) Transformation. Methods Mol Biol 103: 27–39

55. Livak KJ, Schmittgen TD (2001) Analysis of relative gene expression data using real-time quantitative PCR and the 2(-Delta Delta C(T)) Method. Methods 25: 402–8

56. Nguyen T, Bjorkman J, Paton BC, Crane DI (2006) Failure of microtubule-mediated peroxisome division and trafficking in disorders with reduced peroxisome abundance. J Cell Sci 119: 636–45

57. Espeel M, Depreter M, Nardacci R, D’Herde K, Kerckaert I, Stefanini S, Roels F (1997) Biogenesis of peroxisomes in fetal liver. Microsc Res Tech 39: 453–66

58. Mahalingam SS, Shukla N, Farre JC, Zientara-Rytter K, Subramani S (2021) Balancing the Opposing Principles That Govern Peroxisome Homeostasis. Trends Biochem Sci 46: 200–212

59. Reinisch KM, Prinz WA (2021) Mechanisms of nonvesicular lipid transport. J Cell Biol 220

60. Al-Saryi NA, Al-Hejjaj MY, van Roermund CWT, Hulmes GE, Ekal L, Payton C, Wanders RJA, Hettema EH (2017) Two NAD-linked redox shuttles maintain the peroxisomal redox balance in Saccharomyces cerevisiae. Sci Rep 7: 11868

61. Gabay-Maskit S, Cruz-Zaragoza LD, Shai N, Eisenstein M, Bibi C, Cohen N, Hansen T, Yifrach E, Harpaz N, Belostotsky R, et al. (2020) A piggybacking mechanism enables peroxisomal localization of the glyoxylate cycle enzyme Mdh2 in yeast. J Cell Sci 133

62. Kumar S, Singh R, Williams CP, van der Klei IJ (2016) Stress exposure results in increased peroxisomal levels of yeast Pnc1 and Gpd1, which are imported via a piggy-backing mechanism. Biochim Biophys Acta 1863: 148–56

63. Ansell R, Granath K, Hohmann S, Thevelein JM, Adler L (1997) The two isoenzymes for yeast NAD+-dependent glycerol 3-phosphate dehydrogenase encoded by GPD1 and GPD2 have distinct roles in osmoadaptation and redox regulation. EMBO J 16: 2179–87

64. Farre JC, Li P, Subramani S (2021) BiFC Method Based on Intraorganellar Protein Crowding Detects Oleate-Dependent Peroxisomal Targeting of Pichia pastoris Malate Dehydrogenase. Int J Mol Sci 22

65. Lasserre JP, Dautant A, Aiyar RS, Kucharczyk R, Glatigny A, Tribouillard-Tanvier D, Rytka J, Blondel M, Skoczen N, Reynier P, et al. (2015) Yeast as a system for modeling mitochondrial disease mechanisms and discovering therapies. Dis Model Mech 8: 509–26

66. Pinchot GB (1967) The mechanism of uncoupling of oxidative phosphorylation by 2,4-dinitrophenol. J Biol Chem 242: 4577–83

67. Hedbacker K, Carlson M (2008) SNF1/AMPK pathways in yeast. Front Biosci 13: 2408–20

68. Kayikci O, Nielsen J (2015) Glucose repression in Saccharomyces cerevisiae. FEMS Yeast Res 15

69. Kim JH, Roy A, Jouandot D2nd, Cho KH (2013) The glucose signaling network in yeast. Biochim Biophys Acta 1830: 5204-10

70. Saleem RA, Knoblach B, Mast FD, Smith JJ, Boyle J, Dobson CM, Long-O’Donnell R, Rachubinski RA, Aitchison JD (2008) Genome-wide analysis of signaling networks regulating fatty acid-induced gene expression and organelle biogenesis. J Cell Biol 181: 281–92

71. Shi L, Wang X, Wang J, Zhang P, Qi F, Cai M, Zhang Y, Zhou X (2018) Transcriptome analysis of Deltamig1 Deltamig2 mutant reveals their roles in methanol catabolism, peroxisome biogenesis and autophagy in methylotrophic yeast Pichia pastoris. Genes Genomics 40: 399–412

72. Vogl T, Sturmberger L, Fauland PC, Hyden P, Fischer JE, Schmid C, Thallinger GG, Geier M, Glieder A (2018) Methanol independent induction in Pichia pastoris by simple derepressed overexpression of single transcription factors. Biotechnol Bioeng 115: 1037–1050

73. Wang X, Wang Q, Wang J, Bai P, Shi L, Shen W, Zhou M, Zhou X, Zhang Y, Cai M (2016) Mit1 Transcription Factor Mediates Methanol Signaling and Regulates the Alcohol Oxidase 1 (AOX1) Promoter in Pichia pastoris. J Biol Chem 291: 6245–61

74. Lin-Cereghino GP, Godfrey L, de la Cruz BJ, Johnson S, Khuongsathiene S, Tolstorukov I, Yan M, Lin-Cereghino J, Veenhuis M, Subramani S, et al. (2006) Mxr1p, a key regulator of the methanol utilization pathway and peroxisomal genes in Pichia pastoris. Mol Cell Biol 26: 883–97

75. Hou C, Yang Y, Xing Y, Zhan C, Liu G, Liu X, Liu C, Zhan J, Xu D, Bai Z (2020) Targeted editing of transcriptional activator MXR1 on the Pichia pastoris genome using CRISPR/Cas9 technology. Yeast 37: 305–312

76. Wang X, Cai M, Shi L, Wang Q, Zhu J, Wang J, Zhou M, Zhou X, Zhang Y (2016) PpNrg1 is a transcriptional repressor for glucose and glycerol repression of AOX1 promoter in methylotrophic yeast Pichia pastoris. Biotechnol Lett 38: 291–8

77. Shen W, Kong C, Xue Y, Liu Y, Cai M, Zhang Y, Jiang T, Zhou X, Zhou M (2016) Kinase Screening in Pichia pastoris Identified Promising Targets Involved in Cell Growth and Alcohol Oxidase 1 Promoter (PAOX1) Regulation. PLoS One 11: e0167766

78. Sahu U, Krishna Rao K, Rangarajan PN (2014) Trm1p, a Zn(II)(2)Cys(6)-type transcription factor, is essential for the transcriptional activation of genes of methanol utilization pathway, in Pichia pastoris. Biochem Biophys Res Commun 451: 158–64

79. Yan Y, Zhou XE, Xu HE, Melcher K (2018) Structure and Physiological Regulation of AMPK. Int J Mol Sci 19

80. Parua PK, Ryan PM, Trang K, Young ET (2012) Pichia pastoris 14-3-3 regulates transcriptional activity of the methanol inducible transcription factor Mxr1 by direct interaction. Mol Microbiol 85: 282–98

81. Cregg JM, Cereghino JL, Shi J, Higgins DR (2000) Recombinant protein expression in Pichia pastoris. Mol Biotechnol 16: 23–52

82. Hartner FS, Glieder A (2006) Regulation of methanol utilisation pathway genes in yeasts. Microb Cell Fact 5: 39

83. Mizuno T, Masuda Y, Irie K (2015) The Saccharomyces cerevisiae AMPK, Snf1, Negatively Regulates the Hog1 MAPK Pathway in ER Stress Response. PLoS Genet 11: e1005491

84. Vyas VK, Kuchin S, Carlson M (2001) Interaction of the repressors Nrg1 and Nrg2 with the Snf1 protein kinase in Saccharomyces cerevisiae. Genetics 158: 563–72

85. Wang J, Wang X, Shi L, Qi F, Zhang P, Zhang Y, Zhou X, Song Z, Cai M (2017) Methanol-Independent Protein Expression by AOX1 Promoter with trans-Acting Elements Engineering and Glucose-Glycerol-Shift Induction in Pichia pastoris. Sci Rep 7: 41850

86. Farre JC, Mahalingam SS, Proietto M, Subramani S (2019) Peroxisome biogenesis, membrane contact sites, and quality control. EMBO Rep 20

87. Castro IG, Schuldiner M, Zalckvar E (2018) Mind the Organelle Gap - Peroxisome Contact Sites in Disease. Trends Biochem Sci 43: 199–210

88. Chu BB, Liao YC, Qi W, Xie C, Du X, Wang J, Yang H, Miao HH, Li BL, Song BL (2015) Cholesterol transport through lysosome-peroxisome membrane contacts. Cell 161: 291–306

89. Schuldiner M, Zalckvar E (2017) Incredibly close-A newly identified peroxisome-ER contact site in humans. J Cell Biol 216: 287–289

90. Shai N, Schuldiner M, Zalckvar E (2016) No peroxisome is an island - Peroxisome contact sites. Biochim Biophys Acta 1863: 1061–9

91. Diogo CV, Yambire KF, Fernandez Mosquera L, Branco FT, Raimundo N (2018) Mitochondrial adventures at the organelle society. Biochem Biophys Res Commun 500: 87–93

92. Kurihara T, Ueda M, Okada H, Kamasawa N, Naito N, Osumi M, Tanaka A (1992) Beta-oxidation of butyrate, the short-chain-length fatty acid, occurs in peroxisomes in the yeast Candida tropicalis. J Biochem 111: 783–7

93. Tanaka A, Osumi M, Fukui S (1982) Peroxisomes of alkane-grown yeast: fundamental and practical aspects. Ann N Y Acad Sci 386: 183–99

94. Passarella S, de Bari L, Valenti D, Pizzuto R, Paventi G, Atlante A (2008) Mitochondria and L-lactate metabolism. FEBS Lett 582: 3569–76

95. Lanz MC, Yugandhar K, Gupta S, Sanford EJ, Faca VM, Vega S, Joiner AMN, Fromme JC, Yu H, Smolka MB (2021) In-depth and 3-dimensional exploration of the budding yeast phosphoproteome. EMBO Rep 22: e51121

96. Shevade A, Strogolova V, Orlova M, Yeo CT, Kuchin S (2018) Mitochondrial Voltage-Dependent Anion Channel Protein Por1 Positively Regulates the Nuclear Localization of Saccharomyces cerevisiae AMP-Activated Protein Kinase. mSphere 3

97. Hedbacker K, Carlson M (2006) Regulation of the nucleocytoplasmic distribution of Snf1-Gal83 protein kinase. Eukaryot Cell 5: 1950–6

98. Dubois V, Eeckhoute J, Lefebvre P, Staels B (2017) Distinct but complementary contributions of PPAR isotypes to energy homeostasis. J Clin Invest 127: 1202–1214

99. Vamecq J, Papegay B, Nuyens V, Boogaerts J, Leo O, Kruys V (2020) Mitochondrial dysfunction, AMPK activation and peroxisomal metabolism: A coherent scenario for non-canonical 3-methylglutaconic acidurias. Biochimie 168: 53–82

100. Tam YY, Torres-Guzman JC, Vizeacoumar FJ, Smith JJ, Marelli M, Aitchison JD, Rachubinski RA (2003) Pex11-related proteins in peroxisome dynamics: a role for the novel peroxin Pex27p in controlling peroxisome size and number in Saccharomyces cerevisiae. Mol Biol Cell 14: 4089–102

101. Thoms S, Erdmann R (2005) Dynamin-related proteins and Pex11 proteins in peroxisome division and proliferation. FEBS J 272: 5169–81

102. Rinaldi MA, Patel AB, Park J, Lee K, Strader LC, Bartel B (2016) The Roles of beta-Oxidation and Cofactor Homeostasis in Peroxisome Distribution and Function in Arabidopsis thaliana. Genetics 204: 1089–1115

103. Liu S, Liu S, He B, Li L, Li L, Wang J, Cai T, Chen S, Jiang H (2021) OXPHOS deficiency activates global adaptation pathways to maintain mitochondrial membrane potential. EMBO Rep 22: e51606

104. Bergman O, Ben-Shachar D (2016) Mitochondrial Oxidative Phosphorylation System (OXPHOS) Deficits in Schizophrenia: Possible Interactions with Cellular Processes. Can J Psychiatry 61: 457–69

105. Lopez-Gallardo E, Iceta R, Iglesias E, Montoya J, Ruiz-Pesini E (2011) OXPHOS toxicogenomics and Parkinson’s disease. Mutat Res 728: 98–106

106. Zhu Z, Wang X (2017) Significance of Mitochondria DNA Mutations in Diseases. Adv Exp Med Biol 1038: 219–230

107. Olgun A, Akman S (2007) Mitochondrial DNA-deficient models and aging. Ann N Y Acad Sci 1100: 241–5

108. Koopman WJ, Distelmaier F, Smeitink JA, Willems PH (2013) OXPHOS mutations and neurodegeneration. EMBO J 32: 9–29

109. Herzig S, Shaw RJ (2018) AMPK: guardian of metabolism and mitochondrial homeostasis. Nat Rev Mol Cell Biol 19: 121–135

110. Masters C (1997) Gluconeogenesis and the peroxisome. Mol Cell Biochem 166: 159–68

111. Fourcade S, Lopez-Erauskin J, Ruiz M, Ferrer I, Pujol A (2014) Mitochondrial dysfunction and oxidative damage cooperatively fuel axonal degeneration in X-linked adrenoleukodystrophy. Biochimie 98: 143–9

112. Rao X, Huang X, Zhou Z, Lin X (2013) An improvement of the **2^−△△CT^** method for quantitative real-time polymerase chain reaction data analysis. Biostat Bioinforma Biomath 3: 71–85

