## Supplementary Legends, Figures and Tables for "OXPHOS deficiencies affect peroxisome proliferation by downregulating genes controlled by the SNF1 signaling pathway"

**Supplementary Figure Legends**

Figure S1: **Peroxisome metabolites influence peroxisome size.** (A) Fluorescence microscopy of WT, Δ*pot1* and Δ*aox1* Δ*aox2* cells expressing Pex3-GFP driven by the *PEX3* promoter and BFP-SKL driven by the *GAPDH* promoter. Cells were grown in oleate and methanol for 8 h, respectively. (B) Fluorescence microscopy of WT and Δ*pex5* mutant cells expressing GFP-Pex36 driven by the *PEX36* promoter. Bars: 5 µm

Figure S2: **Peroxisome metabolites influence peroxisome size.** Fluorescence microscopy of WT and NADH-shuttling mutant cells expressing Pex3-GFP driven by the *PEX3* promoter and BFP-SKL driven by the *GAPDH* promoter. Cells were grown in oleate and methanol for 8 h, respectively. Bars: 5 µm.

Figure S3: **Cells treated with the OXPHOS uncoupler, DNP, share same peroxisomal protein expression defects with OXPHOS mutants, without affecting Snf1 phosphorylation.** (A) Western blot for several peroxisomal proteins, as well as the total ($^{T}$) the phosphorylated forms ($^{P}$) of Snf1 in cells treated with and without DNP. Specific bands are indicated with an arrow. (B) Ponceau S staining was used as a loading control.

Figure S4: **PKA inhibition or *HOG1* deletion did not rescue peroxisome proliferation defect of Δ*nugM* mutant cells.** Western blot of Aox1 and Pot1 in WT and Δ*nugM* mutant cells, with and without PKA inhibition or *HOG1* deletion. In *P. pastoris,* PKA consists of a regulatory subunit dimer (Bcy1) and two catalytic subunits (Pka_A, UniProt gene name: PAS_chr1-4_0357; Pka_B, UniProt gene name: PAS_chr3_0964). PKA inhibition was obtained by using a strain with a deletion of PKA_B gene and a point mutation in Pka_A (M219G) that renders the kinase sensitive to the drug, 1NM-PP1, which was added when cultures were shifted to methanol or oleate medium. Ponceau S staining was used as a loading control.

Figure S5: **Gal83 nuclear localization during oleate adaptation is inhibited in Δ*nugM* mutant cells.** Fluorescence microscopy of WT and Δ*nugM* mutant cells expressing Gal83-GFP driven by the *GAL83* promoter and the Sec61-mRFP driven by the *SEC61* promoter and decorating the ER and the perinuclear-ER. Bars: 5 μm.

Figure S6: **The Δ*pex11* cells of *P. pastoris* do not phenocopy the OXPHOS mutant, and can proliferate on methanol.** Peroxisomes are labeled with Pex3-GFP. Bar: 5 μm.

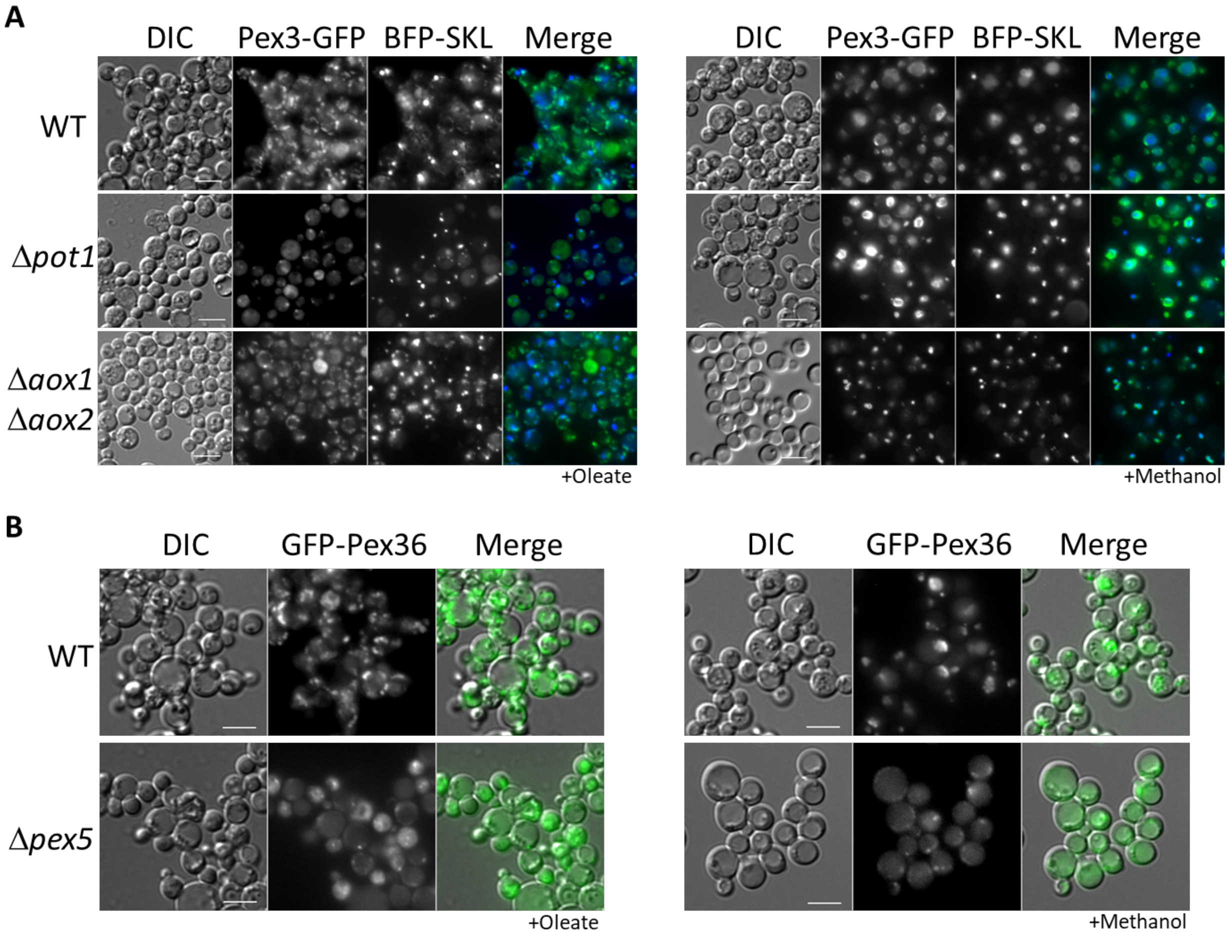
A
DIC
Pex3-GFP
BFP-SKL
Merge
DIC
Pex3-GFP
BFP-SKL
Merge
WT
Δpot1
Δaox1
Δaox2
+Oleate
+Methanol
B
DIC
GFP-Pex36
Merge
DIC
GFP-Pex36
Merge
WT
Δpex5
+Oleate
+Methanol

Figure S1

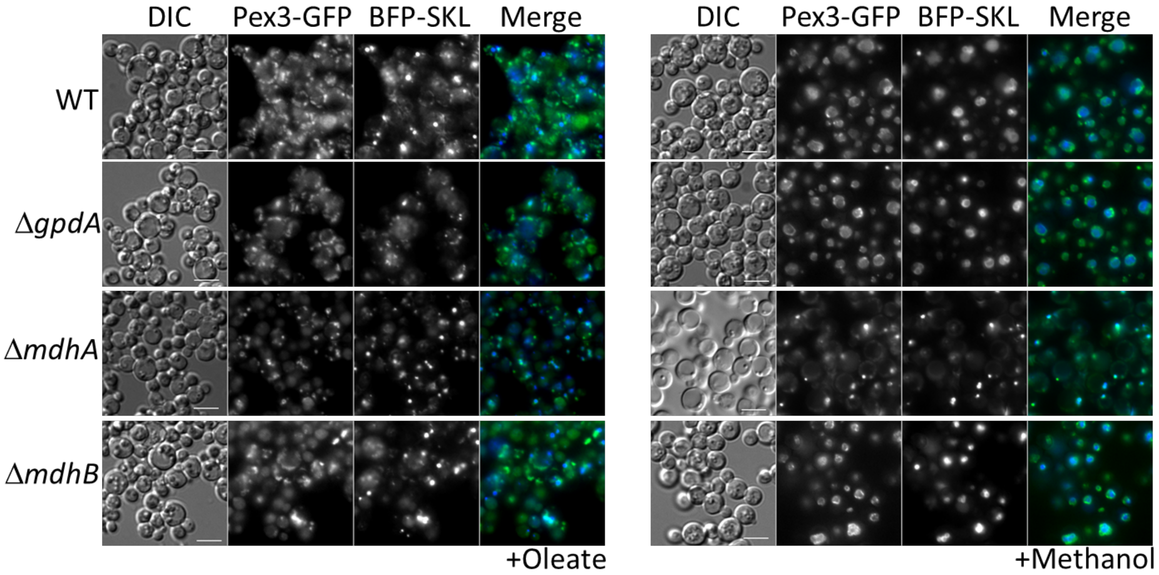
DIC
Pex3-GFP
BFP-SKL
Merge
WT
ΔgpdA
ΔmdhA
ΔmdhB
+Oleate
DIC
Pex3-GFP
BFP-SKL
Merge
+Methanol

Figure S2

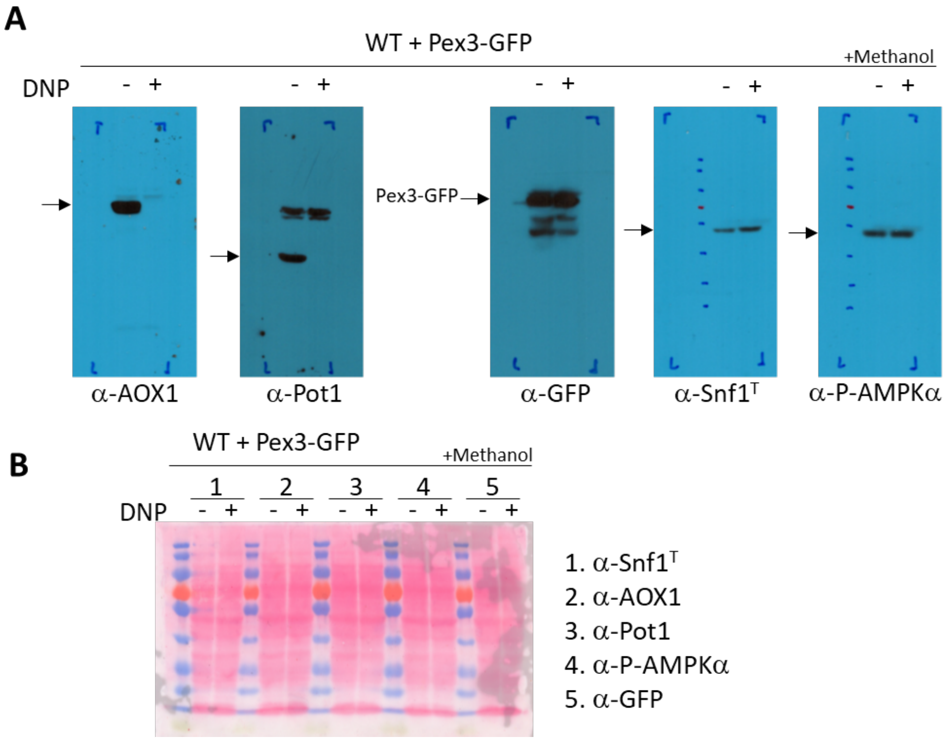
A
WT + Pex3-GFP
+Methanol
DNP
- +
- +
Pex3-GFP
- +
- +
- +
α-AOX1
α-Pot1
α-GFP
α-Snf1T
α-P-AMPKα
B
WT + Pex3-GFP
+Methanol
1
2
3
4
5
DNP
- + - + - + - + - +
1. α-Snf1T
2. α-AOX1
3. α-Pot1
4. α-P-AMPKα
5. α-GFP

Figure S3

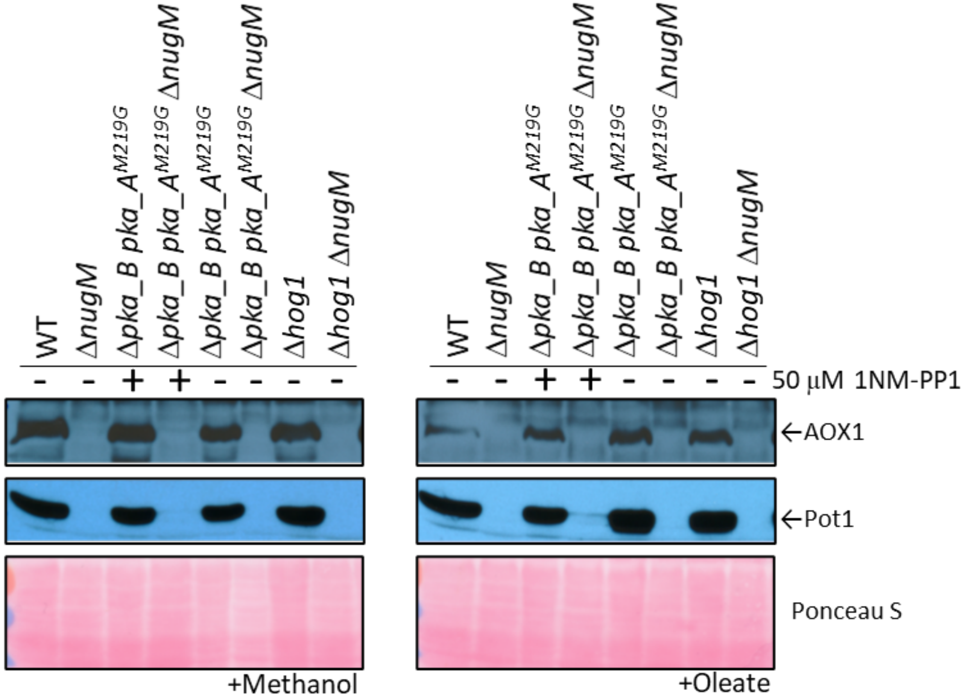
WT
ΔnugM
Δpka_B pka_A^M219G
Δpka_B pka_A^M219G ΔnugM
Δpka_B pka_A^M219G
Δpka_B pka_A^M219G ΔnugM
Δhog1
Δhog1 ΔnugM
- - + + - - - -
- - + + - - - -
50 μM 1NM-PP1
←AOX1
←Pot1
Ponceau S
+Methanol
+Oleate

Figure S4

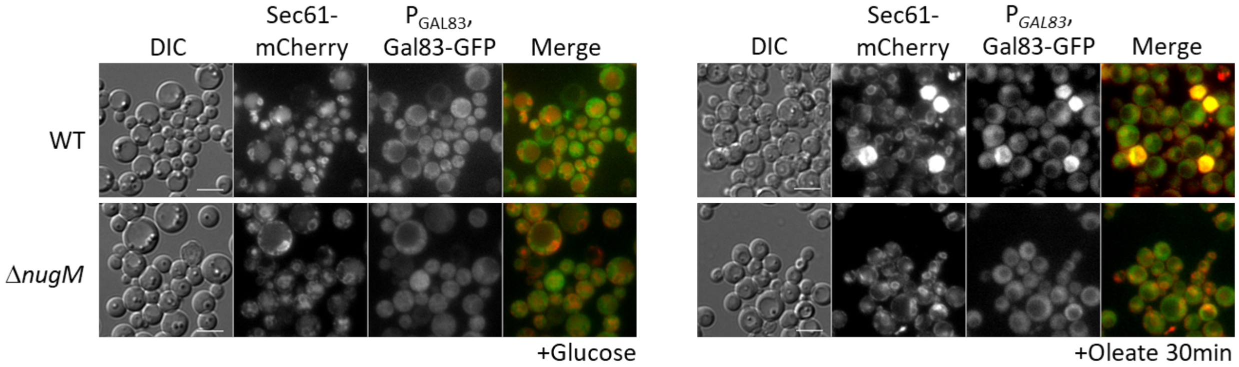
DIC
Sec61-mCherry
$P_{GAL83}$, Gal83-GFP
Merge
WT
ΔnugM
+Glucose
DIC
Sec61-mCherry
$P_{GAL83}$, Gal83-GFP
Merge
+Oleate 30min

Figure S5

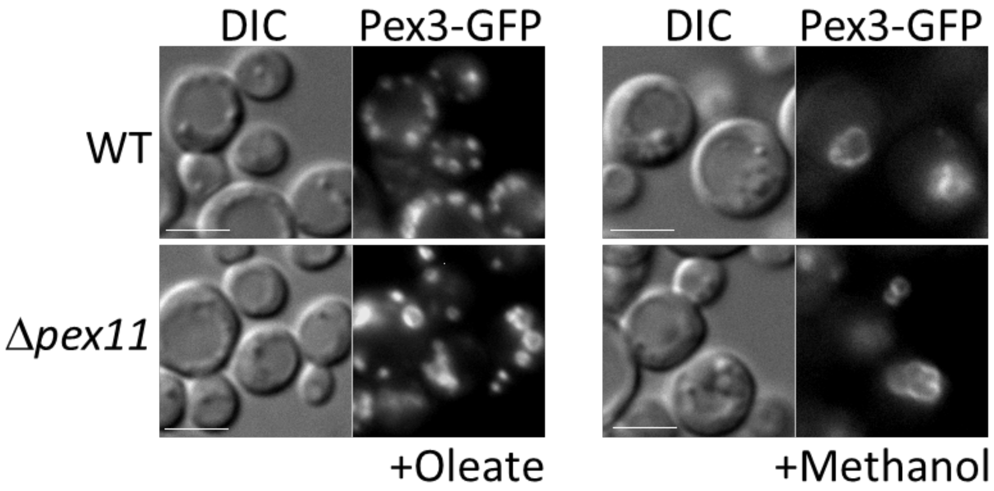


Figure S6

| Name | Stock name | Background | Description | Genotype | References / Sources |
|---|---|---|---|---|---|
| WT | GS115 | NRRL Y-11430 | WT | *his4* | [1] |
| WT | PPY12 | NRRL Y-11430 | WT | *his4 arg4* | [2] |
| WT | PPF1 | NRRL Y-11430 | WT | *his4 arg4* | [3] |
| Δ*gal83* | Δ*PAS_chr1-4_0498* | GS115 | Δ*gal83* | GS115 Δ*gal83*::*ZEO his4* | [4] |
| Δ*sak1* | Δ*PAS_chr2-1_0639* | GS115 | Δ*sak1* | GS115 Δ*sak1*::*ZEO his4* | [4] |
| Δ*tos3?* | Δ*PAS_chr1-3_0213* | GS115 | Δ*tos3?* | GS115 Δ*PAS_chr1-3_0213*::*ZEO his4* | [4] |
| Δ*hog1* | Δ*PAS_chr1-3_0232* | GS115 | Δ*hog1* | GS115 Δ*hog1*::*ZEO his4* | [4] |
| Δ*pka_B* | Δ*PAS_chr3_0964* | GS115 | Δ*pka_B* | GS115 Δ*PAS_chr3_0964*::*ZEO his4* | [4] |
| Δ*mig1* Δ*mig2* Δ*nrg1* | Δ*mig1* Δ*mig2* Δ*nrg1* | GS115 | Δ*mig1* Δ*mig2* Δ*nrg1* | GS115 Δ*mig1*::*ZEO* Δ*mig2*::*KAN* Δ*nrg1*::*HPH his4* | [5] |
| Δ*mxr1* | JC132 | GS115 | *mxr1-1* | GS115 *mxr1-1 his4* | [6] |
| Δ*pex14* | JC404 | NRRL Y-11430 | Δ*pex14* | NRRL Y-11430 Δ*pex14*::*ARG4 his4* | [7] |
| Δ*aox1* Δ*aox2* | MC100-3 | PPF1 | Δ*aox1* Δ*aox2* | PPF1 Δ*aox1*::*ARG4* Δ*aox2*::*HIS4* | [8] |
| Δ*pex5* | *pas8* | PPY12 | Δ*pex5* | PPY12 Δ*pex5*::*ARG4 his4* | [9] |
| Δ*pot1* | Δ*pot1* | PPY12 | Δ*pot1* | PPY12 Δ*pot1*::*ARG4 his4* | [10] |
| WT-OE-Mit1 | WT-Mit1 | GS115 | WT + $P_{GAPDH}$, Mit1 | GS115 $P_{GAPDH}$::pP6GM1($P_{GAPDH}$, Mit1; *HIS4*, *BLA*) | [5] |
| Δ*nugM* | SJCF2769 | GS115 | Δ*nugM* | GS115 Δ*nugM*::*NAT his4* | This study |
| WT + Pex3-GFP + BFP-SKL | SJCF2177 | SEW1 | Δ*pex3* + $P_{PEX3}$, Pex3-GFP + $P_{GAPDH}$, BFP-SKL | PPY12 Δ*pex3*::*ARG4 his4*::pJCF533($P_{PEX3}$, Pex3-GFP; *HIS4*) $P_{GAPDH}$::pJCF742($P_{GAPDH}$, BFP-SKL; *ZEO*) | This study |
| Δ*gpdA* + Pex3-GFP + BFP-SKL | SJCF2649 | SJCF2177 | Δ*gpdA* Δ*pex3* + $P_{PEX3}$, Pex3-GFP + $P_{GAPDH}$, BFP-SKL | PPY12 Δ*gpdA*::*NAT* Δ*pex3*::*ARG4 his4*::pJCF533($P_{PEX3}$, Pex3-GFP; *HIS4*) $P_{GAPDH}$::pJCF742($P_{GAPDH}$, BFP-SKL; *ZEO*) | This study |
| Δ*mdhA* + Pex3-GFP + BFP-SKL | SJCF2650 | SJCF2177 | Δ*mdhA* Δ*pex3* + $P_{PEX3}$, Pex3-GFP + $P_{GAPDH}$, BFP-SKL | PPY12 Δ*mdhA*::*HPH* Δ*pex3*::*ARG4 his4*::pJCF533($P_{PEX3}$, Pex3-GFP; *HIS4*) $P_{GAPDH}$::pJCF742($P_{GAPDH}$, BFP-SKL; *ZEO*) | This study |
| Δ*mdhB* + Pex3-GFP + BFP-SKL | SJCF2651 | SJCF2177 | Δ*mdhB* Δ*pex3* + $P_{PEX3}$, Pex3-GFP + $P_{GAPDH}$, BFP-SKL | PPY12 Δ*mdhB*::*HPH* Δ*pex3*::*ARG4 his4*::pJCF533($P_{PEX3}$, Pex3-GFP; *HIS4*) $P_{GAPDH}$::pJCF742($P_{GAPDH}$, BFP-SKL; *ZEO*) | This study |
| Δ*gpdA* Δ*mdhA* + Pex3-GFP + BFP-SKL | SJCF2652 | SJCF2177 | Δ*gpdA* Δ*mdhA* Δ*pex3* + $P_{PEX3}$, Pex3-GFP + $P_{GAPDH}$, BFP-SKL | PPY12 Δ*gpdA*::*NAT* Δ*mdhA*::*HPH* Δ*pex3*::*ARG4 his4*::pJCF533($P_{PEX3}$, Pex3-GFP; *HIS4*) $P_{GAPDH}$::pJCF742($P_{GAPDH}$, BFP-SKL; *ZEO*) | This study |
| Δ*gpdA* Δ*mdhB* + Pex3-GFP + BFP-SKL | SJCF2653 | SJCF2177 | Δ*gpdA* Δ*mdh*B Δ*pex3* + $P_{PEX3}$, Pex3-GFP + $P_{GAPDH}$, BFP-SKL | PPY12 Δ*gpdA*::*NAT* Δ*mdhB*::*HPH* Δ*pex3*::*ARG4 his4*::pJCF533($P_{PEX3}$, Pex3-GFP; *HIS4*) $P_{GAPDH}$::pJCF742($P_{GAPDH}$, BFP-SKL; *ZEO*) | This study |
| Δ*ndufa9* + Pex3-GFP + BFP-SKL | SJCF2597 | SJCF2177 | Δ*ndufa9* Δ*pex3* + $P_{PEX3}$, Pex3-GFP + $P_{GAPDH}$, BFP-SKL | PPY12 Δ*ndufa9*::*NAT* Δ*pex3*::*ARG4 his4*::pJCF533($P_{PEX3}$, Pex3-GFP; *HIS4*) $P_{GAPDH}$::pJCF742($P_{GAPDH}$, BFP-SKL; *ZEO*) | This study |
| Δ*nugM* + Pex3-GFP + BFP-SKL | SJCF2631 | SJCF2177 | Δ*nugM* Δ*pex3* + $P_{PEX3}$, Pex3-GFP + $P_{GAPDH}$, BFP-SKL | PPY12 Δ*nugM*::*NAT* Δ*pex3*::*ARG4 his4*::pJCF533($P_{PEX3}$, Pex3-GFP; *HIS4*) $P_{GAPDH}$::pJCF742($P_{GAPDH}$, BFP-SKL; *ZEO*) | This study |
| Δ*cyt1* + Pex3-GFP + BFP-SKL | SJCF2645 | SJCF2177 | Δ*cyt1* Δ*pex3* + $P_{PEX3}$, Pex3-GFP + $P_{GAPDH}$, BFP-SKL | PPY12 Δ*cyt1*::*NAT* Δ*pex3*::*ARG4 his4*::pJCF533($P_{PEX3}$, Pex3-GFP; *HIS4*) $P_{GAPDH}$::pJCF742($P_{GAPDH}$, BFP-SKL; *ZEO*) | This study |
| Δ*aox1* Δ*aox2* + Pex3-GFP + BFP-SKL | SJCF2671 | PPF1 | Δ*aox1* Δ*aox2* $P_{PEX3}$, Pex3-GFP + $P_{GAPDH}$, BFP-SKL | PPF1 Δ*aox1*::*ARG4* Δ*aox2*::*HIS4 HIS4*::pJCF571($P_{PEX3}$, Pex3-GFP; *HIS4*, *ZEO*) *HIS4*::pJCF401($P_{GAPDH}$, BFP-SKL; *HIS4*, *KAN*) | This study |
| Δ*pot1* + Pex3-GFP + BFP-SKL | SJCF2655 | Δ*pot1* | Δ*pot1* + $P_{PEX3}$, Pex3-GFP + $P_{GAPDH}$, BFP-SKL | PPY12 Δ*pot1*::*ARG4 PEX3*::pJCF852($P_{PEX3}$, Pex3-GFP; *HIS4*) $P_{GAPDH}$::pJCF742($P_{GAPDH}$, BFP-SKL; *ZEO*) | This study |
| WT + GFP-Pex36 | SJCF1448 | GS115 | WT + $P_{PEX36}$, GFP-Pex36 | GS115 *his4*::pJCF205($P_{PEX36}$, GFP-Pex36; *HIS4*) | This study |
| Δ*pex5* + GFP-Pex36 | SJCF683 | PPY12 | Δ*pex5* + $P_{PEX36}$, GFP-Pex36 | PPY12 Δ*pex5*::*ARG4 his4*::pJCF205($P_{PEX36}$, GFP-Pex36; *HIS4*) | This study |
| WT + Pex11-2HA | SJCF1355 | GS115 | WT + $P_{PEX11}$, Pex11-2HA | GS115 *his4*::pMY59($P_{PEX11}$, Pex11-2HA; *HIS4*) | This study |
| Δ*nugM* + Pex11-2HA | SJCF2709 | SJCF1355 | Δ*nugM* + $P_{PEX11}$, Pex11-2HA | GS115 Δ*nugM*::*NAT his4*::pMY59($P_{PEX11}$, Pex11-2HA; *HIS4*) | This study |
| Δ*gal83* + Pex11-2HA | SJCF2719 | Δ*PAS_chr1-4_0498* | Δ*gal83* + $P_{PEX11}$, Pex11-2HA | GS115 Δ*gal83*::*ZEO his4*::pMY59($P_{PEX11}$, Pex11-2HA; *HIS4*) | This study |
| Δ*sak1* + Pex11-2HA | SJCF2720 | Δ*PAS_chr2-1_0639* | Δ*sak1* + $P_{PEX11}$, Pex11-2HA | GS115 Δ*sak1*::*ZEO his4*::pMY59($P_{PEX11}$, Pex11-2HA; *HIS4*) | This study |
| Δ*tos3?* + Pex11-2HA | SJCF2718 | Δ*PAS_chr1-3_0213* | Δ*tos3?* + $P_{PEX11}$, Pex11-2HA | GS115 Δ*PAS_chr1-3_0213*::*ZEO his4*::pMY59($P_{PEX11}$, Pex11-2HA; *HIS4*) | This study |
| Δ*mxr1* | Δ*mxr1* + pIB1 | JC132 | *mxr1-1* | GS115 *mxr1-1 his4*::pIB1(empty plasmid; *HIS4*) | This study |
| Δ*mit1* | Δ*mit1* + pIB1 + pJCF214 | SMY288 | Δ*mit1* | PPY12 Δ*mit1*::*ZEO his4*::pIB1(empty plasmid; *HIS4*) *arg4*::pJCF214(empty plasmid; *ARG4*) | This study |
| Δ*pka_B* + Pex3-GFP | SJCF2724 | Δ*PAS_chr3_0964* | Δ*pka_B* + $P_{PEX3}$, Pex3-GFP | GS115 Δ*pka_B*::*ZEO PEX3*::pJCF852($P_{PEX3}$, Pex3-GFP; *HIS4*) | This study |

| | | | | | |
|---|---|---|---|---|---|
| Δ*pka_B pka_A*$^{M219G}$ + Pex3-GFP | SJCF2735 | SJCF2724 | Δ*pka_B pka_A*$^{M219G}$ + $P_{PEX3}$, Pex3-GFP | GS115 *PKA_A*::*pka_A*$^{M219G}$(*KAN*) Δ*PAS_chr3_0964*::*ZEO* *PEX3*::pJCF852($P_{PEX3}$, Pex3-GFP; *HIS4*) | This study |
| Δ*pka_B pka_A*$^{M219G}$ Δ*nugM* + Pex3-GFP | SJCF2736 | SJCF2735 | Δ*nugM* Δ*pka_B pka_A*$^{M219G}$ + $P_{PEX3}$, Pex3-GFP | GS115 Δ*nugM*::*NAT* *PKA_A*::*pka_A*$^{M219G}$(*KAN*) Δ*PAS_chr3_0964*::*ZEO* *PEX3*::pJCF852($P_{PEX3}$, Pex3-GFP; *HIS4*) | This study |
| WT + Pex3-GFP | SJCF2722 | GS115 | WT + $P_{PEX3}$, Pex3-GFP | GS115 *PEX3*::pJCF852($P_{PEX3}$, Pex3-GFP; *HIS4*) | This study |
| Δ*gal83* + Pex3-GFP | SJCF2727 | Δ*PAS_chr1-4_0498* | Δ*gal83* + $P_{PEX3}$, Pex3-GFP | GS115 Δ*gal83*::*ZEO* *PEX3*::pJCF852($P_{PEX3}$, Pex3-GFP; *HIS4*) | This study |
| Δ*hog1* + Pex3-GFP | SJCF2729 | Δ*PAS_chr1-3_0232* | Δ*hog1* + $P_{PEX3}$, Pex3-GFP | GS115 Δ*hog1*::*ZEO* *PEX3*::pJCF852($P_{PEX3}$, Pex3-GFP; *HIS4*) | This study |
| Δ*nugM* + Pex3-GFP | SJCF2730 | SJCF2722 | Δ*nugM* + $P_{PEX3}$, Pex3-GFP | GS115 Δ*nugM*::*NAT* *PEX3*::pJCF852($P_{PEX3}$, Pex3-GFP; *HIS4*) | This study |
| Δ*hog1* Δ*nugM* + Pex3-GFP | SJCF2734 | SJCF2729 | Δ*nugM* Δ*hog1* + $P_{PEX3}$, Pex3-GFP | GS115 Δ*nugM*::*NAT* Δ*hog1*::*ZEO* *PEX3*::pJCF852($P_{PEX3}$, Pex3-GFP; *HIS4*) | This study |
| Δ*mig1* Δ*mig2* Δ*nrg1* + Pex3-GFP | SJCF2777 | Δ*mig1* Δ*mig2* Δ*nrg1* | Δ*mig1* Δ*mig2* Δ*nrg1* + $P_{PEX3}$, Pex3-GFP | GS115 Δ*mig1*::*ZEO* Δ*mig2*::*KAN* Δ*nrg1*::*HPH* *PEX3*::pJCF852($P_{PEX3}$, Pex3-GFP; *HIS4*) | This study |
| Δ*mig1* Δ*mig2* Δ*nrg1* Δ*gal83* + Pex3-GFP | SJCF2778 | SJCF2777 | Δ*mig1* Δ*mig2* Δ*nrg1* Δ*gal83* + $P_{PEX3}$, Pex3-GFP | GS115 Δ*gal83*::*NAT* Δ*mig1*::*ZEO* Δ*mig2*::*KAN* Δ*nrg1*::*HPH* *PEX3*::pJCF852($P_{PEX3}$, Pex3-GFP; *HIS4*) | This study |
| Δ*mig1* Δ*mig2* Δ*nrg1* Δ*nugM* + Pex3-GFP | SJCF2779 | SJCF2777 | Δ*mig1* Δ*mig2* Δ*nrg1* Δ*nugM* + $P_{PEX3}$, Pex3-GFP | GS115 Δ*nugM*::*NAT* Δ*mig1*::*ZEO* Δ*mig2*::*KAN* Δ*nrg1*::*HPH* *PEX3*::pJCF852($P_{PEX3}$, Pex3-GFP; *HIS4*) | This study |
| WT + Pex3-GFP | SJCF1520 | GS115 | WT + $P_{PEX3}$, Pex3-GFP | GS115 *his4*::pJCF533($P_{PEX3}$, Pex3-GFP; *HIS4*) | This study |
| Δ*pex14* + Pex3-GFP | SJCF2672 | JC404 | Δ*pex14* + $P_{PEX3}$, Pex3-GFP | NRRL Y-11430 Δ*pex14*::*ARG4* *his4*::pJCF533($P_{PEX3}$, Pex3-GFP; *HIS4*) | This study |
| Δ*pex11* + Pex3-GFP | SJCF1565 | SJCF1555 | Δ*pex11* + $P_{PEX3}$, Pex3-GFP | GS115 Δ*pex11*::*ZEO* *his4*::pJCF533($P_{PEX3}$, Pex3-GFP; *HIS4*) | This study |
| WT + Gal83-GFP | SJCF2772 | GS115 | WT + $P_{GAL83}$, Gal83-GFP | GS115 *his4*::$P_{GAL83}$, Gal83-GFP(*HIS4*) | This study |
| Δ*nugM* + Gal83-GFP | SJCF2773 | SJCF2772 | Δ*nugM* + $P_{GAL83}$, Gal83-GFP | GS115 Δ*nugM*::*NAT* *his4*::$P_{GAL83}$, Gal83-GFP(*HIS4*) | This study |
| Δ*sak1* + Gal83-GFP | SJCF2774 | Δ*PAS_chr2-1_0639* | Δ*sak1* + $P_{GAL83}$, Gal83-GFP | GS115 Δsak1::*ZEO* *his4*::$P_{GAL83}$, Gal83-GFP(*HIS4*) | This study |
| WT + Gal83-GFP + Sec61-mCherry | SJCF2775 | SJCF2772 | WT + $P_{GAL83}$, Gal83-GFP + $P_{SEC61}$, Sec61-mCherry | GS115 *his4*::$P_{GAL83}$, Gal83-GFP(*HIS4*) *SEC61*::$P_{SEC61}$, Sec61-mCherry(*KAN*) | This study |
| Δ*nugM* + Gal83-GFP + Sec61-mCherry | SJCF2776 | SJCF2773 | Δ*nugM* + $P_{GAL83}$, Gal83-GFP + $P_{SEC61}$, Sec61-mCherry | GS115 Δ*nugM*::*NAT* *his4*::$P_{GAL83}$, Gal83-GFP(*HIS4*) *SEC61*::$P_{SEC61}$, Sec61-mCherry(*KAN*) | This study |
| WT + $P_{MXR1}$, Mxr1$^{S215A}$ | SJCF2750 | GS115 | WT + $P_{MXR1}$, Mxr1$^{S215A}$-HA | GS115 *his4*::$P_{MXR1}$, Mxr1$^{S215A}$-HA(*HIS4*) | This study |
| Δ*gal83* + $P_{MXR1}$, Mxr1$^{S215A}$ | SJCF2751 | Δ*PAS_chr1-4_0498* | Δ*gal83* + $P_{MXR1}$, Mxr1$^{S215A}$-HA | GS115 Δ*gal83*::*ZEO* *his4*::$P_{MXR1}$, Mxr1$^{S215A}$-HA(*HIS4*) | This study |
| Δ*nugM* + $P_{MXR1}$, Mxr1$^{S215A}$ | SJCF2752 | SJCF2750 | Δ*nugM* + $P_{MXR1}$, Mxr1$^{S215A}$-HA | GS115 Δ*nugM*::*NAT* *his4*::$P_{MXR1}$, Mxr1$^{S215A}$-HA(*HIS4*) | This study |
| Δ*nugM* + OE-Mit1 | SJCF2764 | WT-Mit1 | Δ*nugM* + $P_{GAPDH}$, Mit1 | GS115 Δ*nugM*::*NAT* $P_{GAPDH}$::pP6GM1($P_{GAPDH}$, Mit1; *HIS4*, *BLA*) | This study |
| Δ*gal83* + OE-Mit1 | SJCF2758 | WT-Mit1 | Δ*gal83* + $P_{GAPDH}$, Mit1 | GS115 Δ*gal83*::*NAT* $P_{GAPDH}$::pP6GM1($P_{GAPDH}$, Mit1; *HIS4*, *BLA*) | This study |
| WT | GS115 + pIB1 | GS115 | WT | GS115 *his4*::pIB1(empty plasmid; *HIS4*) | This study |
| Δ*gal83* | Dgal83 + pIB1 | Δ*PAS_chr1-4_0498* | Δ*gal83* | GS115 Δ*gal83*::*ZEO* *his4*::pIB1(empty plasmid; *HIS4*) | This study |
| Δ*nugM* | Δ*nugM* + pIB1 | SJCF2722 | Δ*nugM* | GS115 Δ*nugM*::*NAT* *his4*::pIB1(empty plasmid; *HIS4*) | This study |

Table S1.

| **Name** | **Primer** | **Target** |
|---|---|---|
| 18s-rRNA-qF | GAGGATTGACAGGATGAGAGC | 18S ribosomal RNA |
| 18s-rRNA-qR | CAAGGTCTCGTTCGTTATCGC | 18S ribosomal RNA |
| Pex11-qPCR-S | GAACAGGAAGGCTCTGAGATT | *PEX11* |

| | | |
|---|---|---|
| PEX11-qPCR-AS | GGTGACCTTGTCGGTTAGTT | *PEX11* |
| AOX-qPCR-s | TCCAGAGGTTCCATTCACATTAC | *AOX1* |
| AOX-qPCR-as | CTTGTAAGCCCAAACCATAGGA | *AOX1* |
| POT1-qPCR-as | CTTCATCCTGGTCCACAGTAATAG | *POT1* |
| POT1-qPCR-s | GAGGAGATTATTCCCATCCAAGTAG | *POT1* |

Table S2.


1. Cregg JM, Barringer KJ, Hessler AY, Madden KR (1985) *Pichia pastoris* as a host system for transformations. *Mol Cell Biol* **5**: 3376-85
2. Gould SJ, McCollum D, Spong AP, Heyman JA, Subramani S (1992) Development of the yeast *Pichia pastoris* as a model organism for a genetic and molecular analysis of peroxisome assembly. *Yeast* **8**: 613-28
3. Cregg J, Madden KR (1987) *Development of yeast transformation systems and construction of methanol-utilization-defective mutants of Pichia pastoris by gene disruption.* In Biological research on industrial yeasts, Stewart GG, Russell RD, Klein RD, Hiebesch RR (eds) pp 1-18. Boca Raton, Florida: CRC Press, inc.
4. Shen W, Kong C, Xue Y, Liu Y, Cai M, Zhang Y, Jiang T, Zhou X, Zhou M (2016) Kinase Screening in *Pichia pastoris* Identified Promising Targets Involved in Cell Growth and Alcohol Oxidase 1 Promoter (PAOX1) Regulation. *PLoS One* **11**: e0167766
5. Wang J, Wang X, Shi L, Qi F, Zhang P, Zhang Y, Zhou X, Song Z, Cai M (2017) Methanol-Independent Protein Expression by AOX1 Promoter with trans-Acting Elements Engineering and Glucose-Glycerol-Shift Induction in *Pichia pastoris*. *Sci Rep* **7**: 41850
6. Johnson MA, Waterham HR, Ksheminska GP, Fayura LR, Cereghino JL, Stasyk OV, Veenhuis M, Kulachkovsky AR, Sibirny AA, Cregg JM (1999) Positive selection of novel peroxisome biogenesis-defective mutants of the yeast *Pichia pastoris*. *Genetics* **151**: 1379-91
7. Johnson MA, Snyder WB, Cereghino JL, Veenhuis M, Subramani S, Cregg JM (2001) *Pichia pastoris* Pex14p, a phosphorylated peroxisomal membrane protein, is part of a PTS-receptor docking complex and interacts with many peroxins. *Yeast* **18**: 621-41
8. Cregg JM, Madden KR, Barringer KJ, Thill GP, Stillman CA (1989) Functional characterization of the two alcohol oxidase genes from the yeast *Pichia pastoris*. *Mol Cell Biol* **9**: 1316-23
9. McCollum D, Monosov E, Subramani S (1993) The pas8 mutant of *Pichia pastoris* exhibits the peroxisomal protein import deficiencies of Zellweger syndrome cells--the PAS8 protein binds to the COOH-terminal tripeptide peroxisomal targeting signal, and is a member of the TPR protein family. *J Cell Biol* **121**: 761-74
10. Elgersma Y, Elgersma-Hooisma M, Wenzel T, McCaffery JM, Farquhar MG, Subramani S (1998) A mobile PTS2 receptor for peroxisomal protein import in *Pichia pastoris*. *J Cell Biol* **140**: 807-20
